# Analyzing the link between RNA secondary structures and R-loop formation with tree polynomials

**DOI:** 10.1101/2023.09.24.559224

**Authors:** Pengyu Liu, Jacob Lusk, Nataša Jonoska, Mariel Vázquez

## Abstract

R-loops are a class of non-canonical nucleic acid structures that typically form during transcription when the nascent RNA hybridizes the DNA template strand, leaving the DNA coding strand unpaired. Co-transcriptional R-loops are abundant in nature and biologically relevant. Recent research shows that DNA sequence and topology affect R-loops, yet it remains unclear how these and other factors drive R-loop formation. In this work, we investigate a link between the secondary structure of the nascent RNA and the probability of R-loop formation. We introduce tree-polynomial representations, a class of mathematical objects that enable accurate and efficient data analysis of RNA secondary structures. With tree-polynomials, we establish a strong correlation between the secondary structure of the RNA transcript and the probability of R-loop formation. We identify that branches with short stems separated by multiple ‘bubbles’ in the RNA secondary structure are associated with the strong correlation and are predictive of R-loop formation.

## Introduction

R-loops are three-stranded nucleic acid structures abundant in the genomes of bacteria, plants and mammals, and play important roles in physiological and pathological processes [4, 8, 23, 32]. The mechanisms of R-loop formation and the factors that drive their occurrence remain unclear. Studies involving genome mapping, biochemical reconstitution and energy-based mathematical models indicate that DNA sequence and topology strongly influence R-loop formation [4]. In particular, R-loops correlate with GC-rich and CG-skew sequences on the template strand as well as negative supercoiling [6, 29]. Co-transcriptional R-loops form during transcription when the nascent RNA hybridizes with the DNA template leaving the unpaired DNA coding strand free to wrap around the RNA:DNA duplex or to adopt a non-standard structure [2, 3, 22]. Here, we investigate a link between the secondary structure of the co-transcriptonal RNA and the formation of R-loops.

The secondary structure of an RNA molecule can be described by its paired and unpaired nucleotides. The paired nucleotides form double-stranded helices that are interspersed with unpaired nucleotides [16] that form loops or bubbles. We can represent an RNA secondary structure with a graph, a mathematical object consisting of vertices (nodes) connected by edges (Figure 1) [25]. Rooted trees are graphs that provide a simple and efficient way to capture essential features of RNA secondary structures without pseudoknots [25, 27]. The use of rooted trees opens the door to a wealth of efficient graph theoretical methods for designing, comparing and analyzing RNA secondary structures [11, 24, 26]. We characterize four features of RNA secondary structures that rooted trees can record: loop-stem relation, loop size, stem size and loop group. We systematically describe six types of rooted trees that record the loop-stem relation and differ in their ability to capture the loop size, the stem size and the loop group. The six representations include two new rooted trees and the previously studied loop-stem tree [25] and arc tree [27].

**Figure 1:**
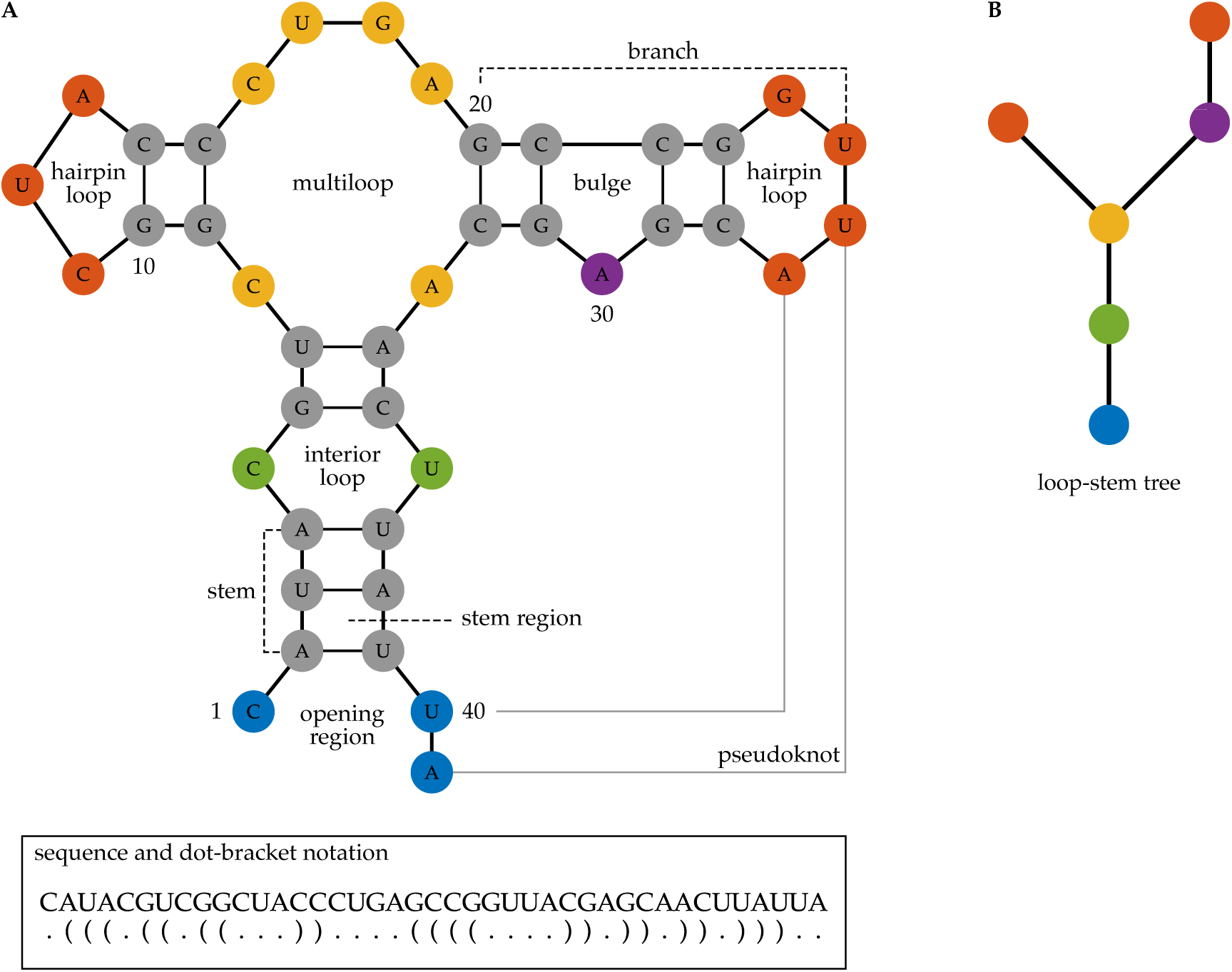
RNA secondary structure, dot-bracket notation and rooted tree representation. Panel A shows the secondary structure of a 41-nucleotide long RNA molecule. Each node represents a nucleotide (A, U, G, or C). The colors indicate different regions of the RNA secondary structure. Integers indicate the nucleotide position along the RNA sequence. Solid black edges represent covalent or hydrogen bonds, and gray edges represent bonds between distant nucleotides forming a pseudoknot. The bottom insert shows the sequence of the RNA molecule and the dot-bracket notation. Panel B shows the loop-stem tree representation of the RNA secondary structure in panel A without the pseudoknot. Here, every vertex represents a loop, and the colors of the vertices correspond to different regions. The blue vertex representing the opening region is the root of the tree. Every edge in the tree represents a stem of the RNA secondary structure.

Structural polynomials are powerful mathematical objects that record important information of discrete structures. Examples include the Tutte polynomial for graphs [30] and the Jones polynomial for knots [12]. Structural polynomials, encoded as coefficient vectors or matrices, have the advantage of being compatible with data analytic tools, including distance-based and machine learning methods. In [13], the author introduced a method that assigns every unlabeled tree a unique 2-variable polynomial representation; we refer to it as *polynomial P*. Recent studies have used the polynomial *P* to analyze pathogen evolution [14] and the syntax of languages [15].

First, we apply the polynomial *P* to rooted tree representations of RNA for studying RNA secondary structures. To describe all information captured by the newly introduced rooted tree RNA representations, we generalize the polynomial *P* to a 3-variable *polynomial Q*. To each of the six tree RNA representations, we define six *tree-polynomials* for an RNA secondary structure (Figure 2). These tree-polynomials allow for accurate, comprehensive, interpretable and easy-to-compute data analysis of RNA secondary structures. We validate our approach by applying tree-polynomials to cluster secondary structures of non-coding RNAs (ncRNAs) obtained from the bpRNA-1m database [5] and show that tree-polynomials can distinguish different families of ncRNA secondary structures.

**Figure 2:**
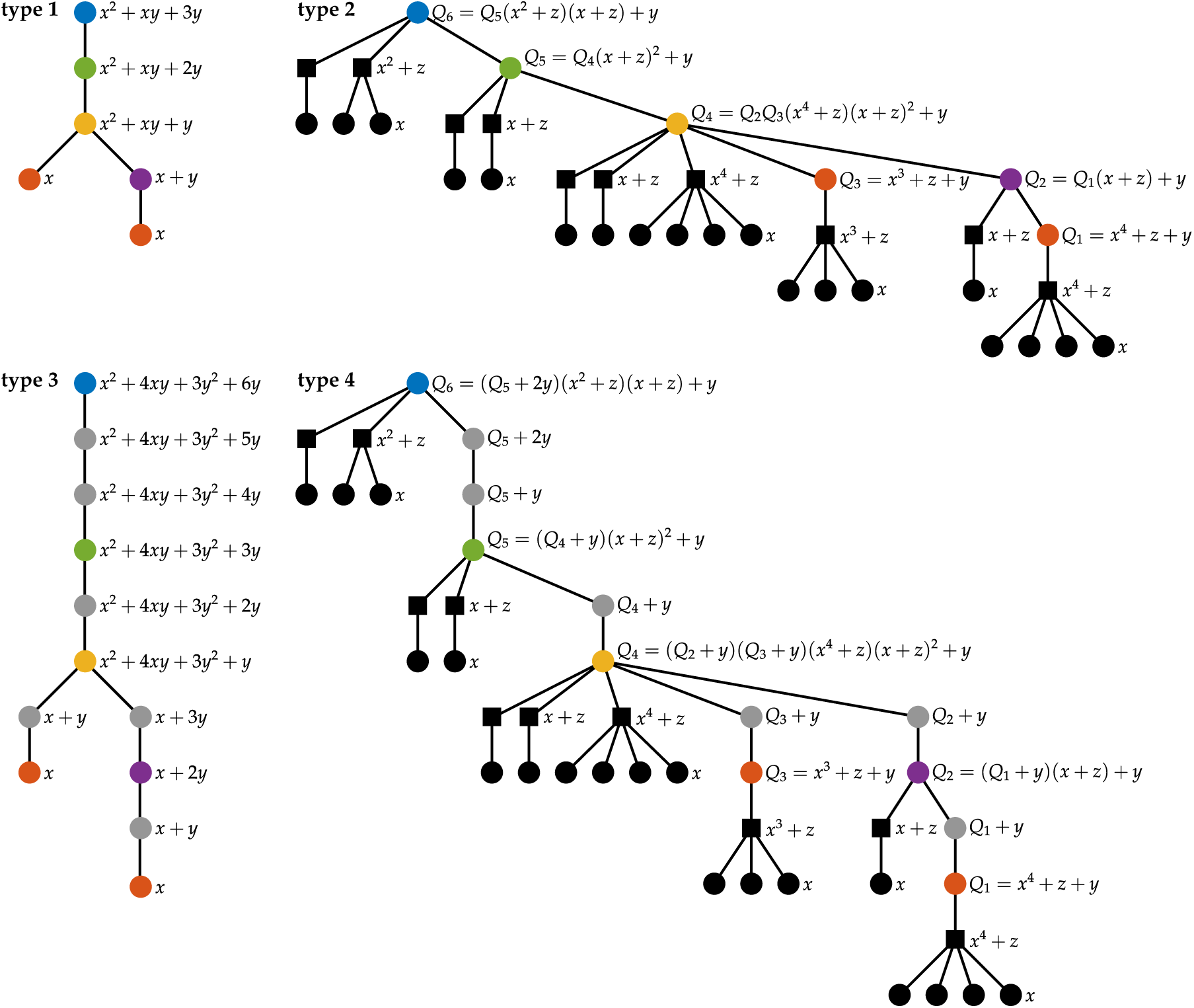
Tree-polynomial representations of an RNA secondary structure. The figure shows the first four types of tree-polynomial representations of the RNA secondary structure shown in Figure 1. Each tree-polynomial representation of an RNA secondary structure consists of a rooted tree representation and its corresponding polynomial. Vertices in the trees have the same colors as the loops and stem regions that they represent. The black round vertices represent unpaired nucleotides. The black square vertices in the type 2 and the type 4 tree representations indicate artificial internal vertices introduced to group unpaired nucleotides. In each rooted tree representation, we illustrate the recursive process of computing the corresponding polynomial from leaf vertices to the root vertex. The polynomial at the root vertex (blue) of a tree is the polynomial that represents the tree.

Second, we use tree-polynomials to analyze a link between the co-transcriptional secondary structure of the RNA and the formation of R-loops. We use the secondary structures of the nascent RNA obtained by DrTransformer, a co-transcriptional RNA folding model [1]. To each of the acquired secondary structures, we associate the six tree-polynomials and compute their coefficient sums. We show that the coefficient sums highly correlate with the experimentally observed probabilities of R-loop formation. We characterize that RNA secondary structures with large coefficient sums have many linear branches with short stems separated by multiple bulges and interior loops. These structural features potentially increase the chance of the nascent RNA hybridizing with the DNA template strand, hence the probability of R-loop formation.

## Results

### Tree-polynomial representations of RNA secondary structures

#### Four features of RNA secondary structures and tree representations

Rooted trees can record four structural features of an RNA secondary structure. The *loop-stem relation* describes how the loops and stems are connected. The *loop size* describes the number of unpaired nucleotides around each loop. The *stem size* describes the number of base pairs in each stem. The *loop group* describes how the stems around a loop separate the unpaired nucleotides into groups. See Methods for more details.

We consider six different rooted tree representations of RNA secondary structures based on the structural features that they record. Table 1 summarizes the features captured by each tree. All the trees take into account the loop-stem relation. *Type 1* trees are the loop stem trees [25]; they only record the loop-stem relation. *Type 2* trees record the loop-stem relation, the loop size and the loop group. *Type 3* trees record the loop-stem relation and the stem size. *Type 4* trees record all four features. For example, see Figure 2. *Type 5* trees record the loop-stem relation and the loop size, and *type 6* trees record the loop-stem relation, the loop size and the stem size. By definition, type 6 trees are the same as arc trees [27]. See Supplementary Figure 1 for an example. The newly defined type 2 and the type 4 trees that record the loop group and the loop size can differentiate RNA secondary structures that are indistinguishable by the other four trees (Supplementary Figure 2). See Supplementary material for more details on how to construct the tree representations from an RNA secondary structure.

**Table 1:** Properties and performance of the six tree-polynomial representations of RNA secondary structures. The first two columns show the types of tree representations and the corresponding polynomials. Obelisks (†) indicate previously studied tree representations in [25, 27]. The middle four columns show the features (loop-stem relation, loop size, stem size and loop group) captured by each tree-polynomial representation. The last three columns show the mean misclassification rates (MMR) in clustering the 735 ncRNA secondary structures in the bpRNA-Rfam dataset (see Methods) and the Pearson’s correlation coefficients (PCC) between the scaled sums (with the highest PCC in the last 10 transcription steps) and the R-loop formation probabilities of linearized supercoiled pFC8 and pFC53 plasmid respectively. Asterisks indicate the top performing tree-polynomial representations in each of these experiments.

| Tree type | Poly-nomial | Loop-stem | Loop size | Stem size | Loop group | MMR | PCC pFC8 | PCC pFC53 |
| --- | --- | --- | --- | --- | --- | --- | --- | --- |
| 1<sup>†</sup> | P | ✓ |  |  |  | 6.82% | 0.77 | 0.79 |
| 2 | Q | ✓ | ✓ |  | ✓ | 17.07% | 0.96 | 0.96 |
| 3 | P | ✓ |  | ✓ |  | 3.40%* | -0.30 | -0.19 |
| 4 | Q | ✓ | ✓ | ✓ | ✓ | 17.51% | 0.97* | 0.97* |
| 5 | P | ✓ | ✓ |  |  | 9.29% | 0.60 | 0.64 |
| 6<sup>†</sup> | P | ✓ | ✓ | ✓ |  | 9.70% | -0.31 | -0.21 |

#### Tree-polynomial representations of RNA secondary structures

Polynomial *P* is a two-variable polynomial that characterizes rooted tree representations of RNA secondary structures [13]. See Methods for more details. We apply polynomial *P* to type 1, 3, 5 and 6 trees. Figure 2 illustrates the recursive process used to compute polynomial *P* on type 1 and type 3 trees.

We generalize the polynomial *P* to *polynomial Q*, a tree distinguishing polynomial with three variables, and apply polynomial *Q* to the new type 2 and type 4 trees to better describe the loop group feature that they record. Specifically, polynomial *Q* differentiates between the artificial internal vertices introduced to group unpaired nucleotides and the internal vertices that represent loops or stem regions in type 2 and type 4 trees. Let *T* be the type 2 (or type 4) tree representation of an RNA secondary structure. We define the polynomial *Q*(*T*, *x*, *y*, *z*) following a recursive process from the leaf vertices to the root vertex. Let *v* be a vertex in the tree *T*. If *v* is a leaf vertex, then *Q*(*v*, *x*, *y*, *z*) = *x*. If *v* is an artificial internal vertex with *k* child vertices *u*_1_, *u*_2_, …, *u_k_*, then *Q*(*v*, *x*, *y*, *z*) = *z* + Π*^k^_*i = 1*_ Q*(*u_i_*, *x*, *y*, *z*). If *v* is an internal vertex that represents a loop or a stem region and has *k* child vertices *u*_1_, *u*_2_, …, *u_k_*, then *Q*(*v*, *x*, *y*, *z*) = *y* + Π*^k^_*i = 1*_ Q*(*u_i_*, *x*, *y*, *z*). The polynomial *Q* for the entire tree *T* is defined as the polynomial at the root vertex of the tree. The variables *x*, *y* and *z* encode structural information of the unpaired nucleotides, the loops and the groups of unpaired nucleotides around each loop, respectively. Figure 2 illustrates the recursive process used to compute polynomial *Q* on type 2 and type 4 tree representations. Note that substituting the variable *z* with *y* in the polynomial *Q*(*T*, *x*, *y*, *z*) of a type 2 (or type 4) tree *T* yields the polynomial *P*(*T*, *x*, *y*) for the tree *T*, which does not distinguish the artificial internal vertices in *T*.

We define the *tree-polynomial representation*, or *tree-polynomial* for short, of a given RNA secondary structure as the pair consisting of a tree representation and its corresponding polynomial, as in Table 1. The name of a tree-polynomial corresponds to the name of the tree used to define it. For example, the *type 1 tree-polynomial* is the pair consisting of the type 1 tree and polynomial *P*.

#### Tree-polynomial distances for RNA secondary structure comparison

We measure the similarity between RNA secondary structures using distance-based methods. The *polynomial P distance*, defined in Methods, is the distance between two rooted trees based on polynomial *P*. We generalize the polynomial *P* distance for polynomial *Q* by representing the polynomial *Q*(*T*, *x*, *y*, *z*) of degree *n* for a rooted tree *T* as an (*n* + 1) *×* (*n* + 1) *×* (*n* + 1) coefficient matrix. Each entry of the coefficient matrix *c*^(^*^i^*^,*j*,*k*)^ is the coefficient of the term *c*^(^*^i^*^,*j*,*k*)^ *x^i^y^j^z^k^* in *Q*(*T*, *x*, *y*, *z*), where 0 *≤ i*, *j*, *k ≤ n*. Given two type 2 (or type 4) trees *S* and *T*, let *b*^(^*^i^*^,*j*,*k*)^ and *c*^(^*^i^*^,*j*,*k*)^ be the coefficients in *Q*(*S*, *x*, *y*, *z*) and *Q*(*T*, *x*, *y*, *z*), respectively. We define the *polynomial Q distance* between *S* and *T*, *d_Q_*(*S*, *T*), as in Formula (1), where the function *κ* is defined by Formula (2).

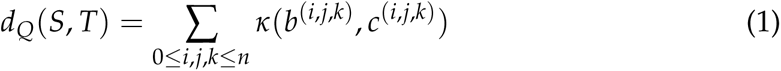

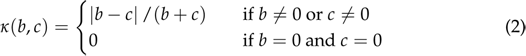

#### Tree-polynomial distances differentiate non-coding RNA families

To validate our approach, we apply the six tree-polynomials and their corresponding polynomial distances to cluster the 735 secondary structures of non-coding RNAs (ncRNAs) in the bpRNA-Rfam dataset [5, 7]. The bpRNA-Rfam dataset contains secondary structures from seven families of ncRNAs including 5.8S ribosomal RNA and U12 minor spliceosomal RNA that play an important role in gene regulation and editing [9, 19]. The ncRNAs in the dataset have similar length and distinct secondary structures. See Methods and Supplementary material for more information about the bpRNA-Rfam dataset.

First, we cluster the ncRNA secondary structures in the bpRNA-Rfam dataset by applying the k-medoids clustering algorithm [28] to the pairwise tree-polynomial distances of the secondary structures. Then, we compare the clustering results to the true families of the ncRNAs. In Table 1, we report the mean misclassification rate (MMR) for each of the six tree-polynomials and their corresponding polynomial distance. The MMRs measures the clustering accuracy. The type 3 tree-polynomial has the highest accuracy in clustering ncRNA secondary structures in the bpRNA-Rfam dataset (MMR = 3.40%). The type 1 tree-polynomial has the second top performance (MMR = 6.82%). Supplementary Figures 4-5 illustrate the pairwise tree-polynomial distances based on each type of tree-polynomial.

The clustering results for the ncRNA secondary structures in the bpRNA-Rfam dataset suggest that tree-polynomials that record more features do not necessarily perform better than those recording fewer features: Type 1 and 3 tree-polynomials perform better than type 2 and 4. This is partially because ncRNA secondary structures in the same family but from different organisms have similar loop-stem relations and stem sizes but largely different loop sizes. See Supplementary Figure 6-7 for examples ncRNA secondary structures in the bpRNA-Rfam dataset.

### The link between RNA secondary structure and R-loop formation

#### The nascent RNA secondary structure correlates with R-loop formation

R-loops typically form behind the polymerase soon after transcription when the RNA hybridizes with the template DNA. We hypothesize that there is a link between the secondary structure of the nascent RNA and the probability of R-loop formation. Tree polynomials allow us to test this hypothesis. We focus on two plasmid sequences that carry mammalian genes and are known to form R-loops, pFC8 and pFC53 [29]. First, we scan each plasmid’s amplicon region with a 200nt-long sliding window. For each 200nt-long RNA transcript, we predict its secondary structure at each transcription step using DrTransformer, an energy-based co-transcriptional RNA folding model [1]. We compute the sum of coefficients of the type *k* tree-polynomial representation (*k* = 1, 2, …, 6) of each predicted secondary structure and call it the *type k coefficient sum* of the secondary structure. In this way, to each predicted RNA secondary structure correspond six coefficient sums. Next, we compare the coefficient sums at a transcription step to the probability of R-loop formation at that nucleotide. For each *k*, the *type k scaled sum* of a plasmid is based on the type *k* coefficient sums at a transcription step in a way that imitates the construction of the experimental probability of R-loop formation [17, 18, 29]. See Methods and Supplementary Figure 8-9 for more details about the plasmids, the definitions and the process. The scaled sums of RNA secondary structures at the last 10 transcription steps provide structural information for majority of the 200nt RNA strand; we focus on these structures. See the Supplementary material (Supplementary Table 3-4) and the interactive 3D figures available at https://pliumath.github.io/rloops.html for analogous results at other transcription steps.

We observe a strong positive correlation between the probabilities of R-loop formation and the types 1, 2 and 4 scaled sums of pFC8 and pFC53, see Figure 3 and Supplementary Figures 10-11. The corresponding Pearson’s correlation coefficients (PCCs) confirm the strong correlation, see Table 1 and Supplementary Table 3-4. In [29] the authors identified four R-loop forming sequence clusters in the amplicon region of pFC8 and three in the amplicon region of pFC53. For each plasmid, the *major peak* refers to the sequence cluster with the highest probability of R-loop formation; similarly, the sequence cluster with the second highest probability of R-loop formation is the *minor peak* (Supplementary Figure 10). The type 2 and the type 4 scaled sums closely match the probabilities of R-loop formation at the major peak with high PCCs (*≥* 0.96) (Table 1 and Figure 3). Despite relatively lower PCCs (*≥* 0.77), the type 1 scaled sums suggest the positions of other R-loop forming sequence clusters.

**Figure 3:**
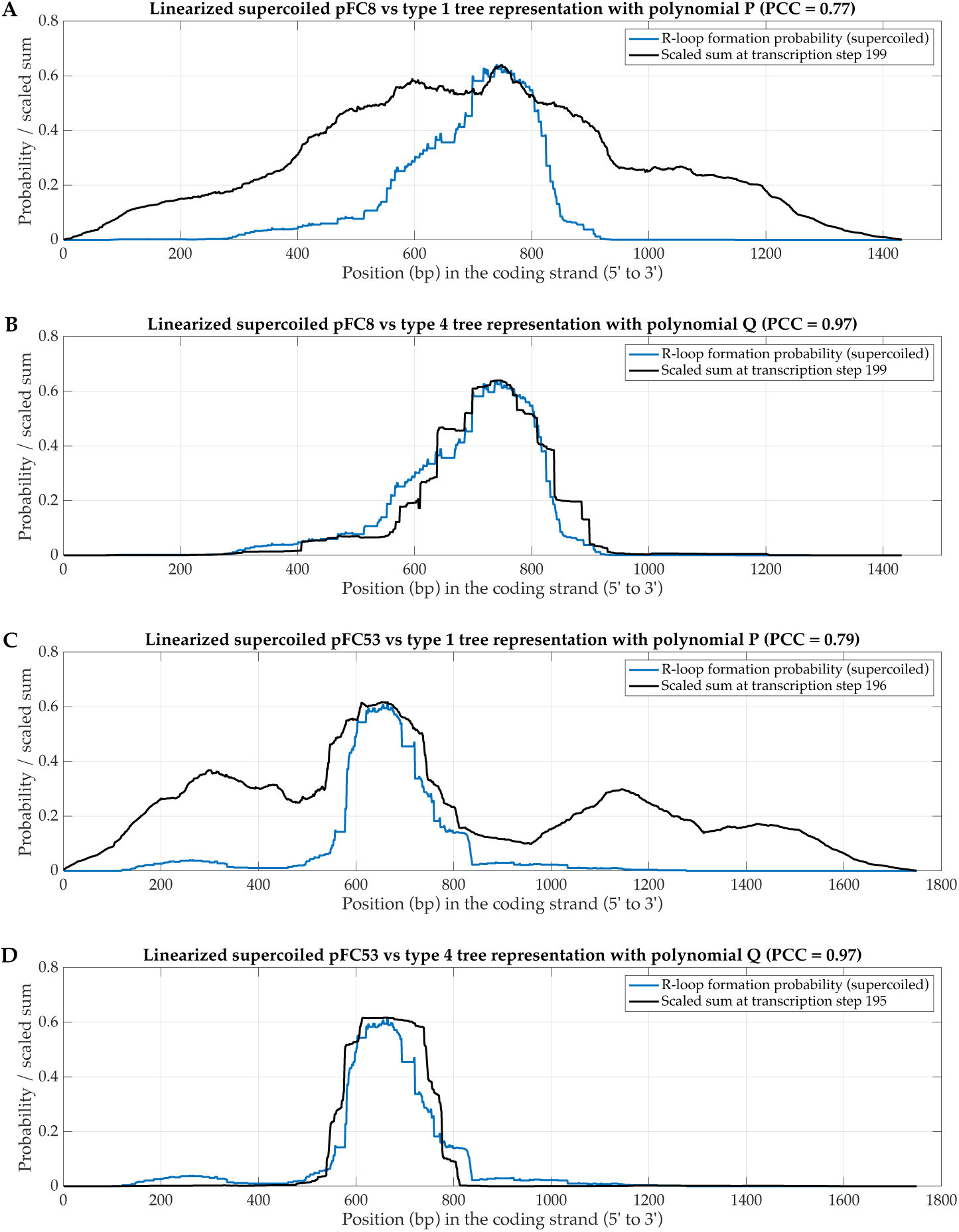
The correlations between the scaled sums and the R-loop formation probabilities of linearized supercoiled pFC8 and pFC53 plasmids. The figure shows the experimental probability of R-loop formation for the linearized supercoiled pFC8 plasmid with the type 1 (panel A) and the type 4 (panel B) scaled sums, and the experimental probability of R-loop formation for the linearized supercoiled pFC53 plasmid with the type 1 (panel C) and the type 4 (panel D) scaled sums. The experimental probabilities of R-loop formation are from [29]. The displayed scaled sums have the highest PCC in the last 10 transcription steps.

#### Branches with bubbles separating short stems drive the strong correlation

Next, we investigate what features of RNA secondary structures contribute to large coefficient sums. Figure 4 shows the structures with the two largest type 1 coefficient sums. Likewise, Supplementary Figures 12-13 show the structures with the two largest type 2 and type 4 coefficient sums, respectively. Both of these structures have linear branches with multiple bulges and interior loops separated by short stems. We refer to the bulges and interior loops in a linear branch as *bubbles*. Note that a linear branch corresponds to a path in the type 1 tree of an RNA secondary structure. The length of a path depends on the number of bubbles in the linear branch. The RNA secondary structures that correspond to large type 1 coefficient sums contain several linear branches with multiple bubbles, as observed in Figure 4. This is because the type 1 tree-polynomial for each linear branch is a path with polynomial *x* + *ly*, where *l* indicates the number of bubbles in the branch. Multiplying the polynomials for all the linear branches to obtain the type 1 tree-polynomial of the secondary structure causes the coefficient sum to increase exponentially with the number of bubbles.

**Figure 4:**
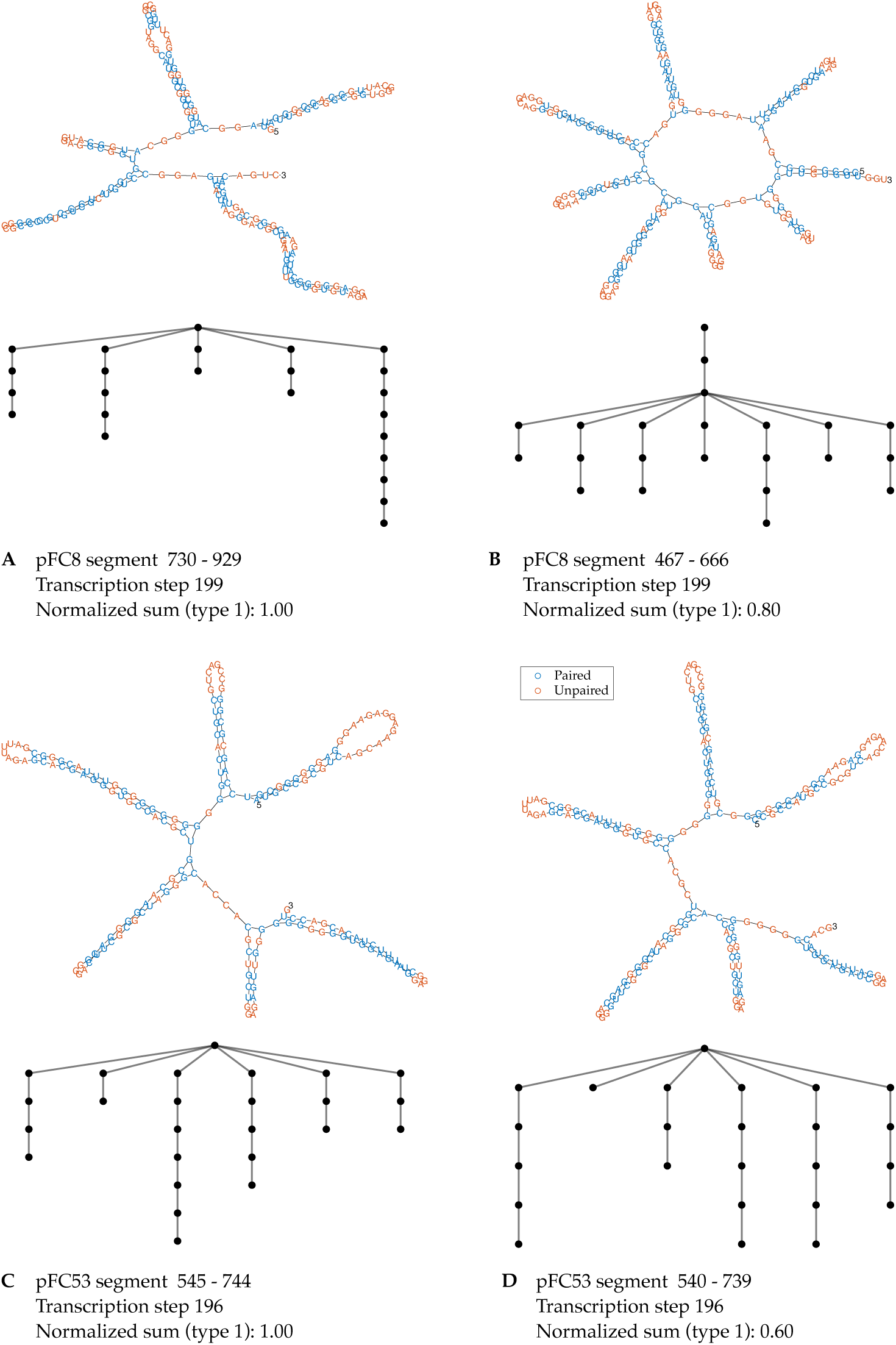
Secondary structures of RNA segments with the largest type 1 coefficient sums. The figure shows the secondary structures of RNA segments of pFC8 (panel A and B) and of pFC53 (panel C and D) with the two largest type 1 coefficient sums at the transcription step with the highest PCC in the last 10 transcription steps. Type 1 tree representations follow the corresponding secondary structures.

Type 1 and type 2 trees of a linear branch contain a path of vertices that represent loops as in Figure 2. The loop-representing vertices in type 2 trees have descendant vertices. Specifically, a vertex in the path representing a bulge has one child vertex indicating the group of unpaired nucleotides, with polynomial *x^k^* + *z*. A vertex in the path representing an interior loop has two child vertices representing the two groups of unpaired nucleotides, with polynomials *x^k^* + *z* and *x^k′^* + *z*. Here, the exponents *k* and *k^′^* indicate the number of unpaired nucleotides in each group. The type 4 tree of a linear branch has the same structure as a type 2 tree with additional vertices corresponding to stem regions. While type 1 trees do not distinguish between bulges and interior loops, interior loops contribute more than bulges to the coefficient sums of the type 2 and the type 4 tree-polynomials of a linear branch. This is because an interior loop provides two polynomial factors *x^k^* + *z* and *x^k′^* + *z*, a bulge provides only one polynomial factor *x^k^* + *z*, and the coefficient sums increase exponentially with respect to the number of such polynomial factors. In sum, the linear branches of RNA secondary structures with large type 2 and type 4 coefficient sums have more interior loops than bulges (Supplementary Figure 12-13).

We now focus on RNA secondary structures near the 3’ end of the amplicon region, where few R-loops form (Supplementary Figure 17). These secondary structures contain few linear branches, and most of the linear branches have longer stems separating fewer bubbles. Compared to the secondary structures in the major peaks shown in Figure 4, the secondary structures in the minor peaks have fewer bubble-rich linear branches, and more bubbles are bulges rather than interior loops (Supplementary Figure 18). This partially explains why the type 2 and type 4 scaled sums do not capture the minor peaks.

Based on these observations, we postulate that the base pairs in bubble-rich linear branches are prone to breaking due to short stems separated by bubbles. Broken base pairs in the secondary structure make it easier for the nascent RNA to pair with the DNA template strand, thus increasing the probability of R-loop formation.

## Discussion

Rooted trees are convenient objects used to represent RNA secondary structures without pseudoknots. In this work, we systematically defined six different rooted tree representations that record combinations of four important features of RNA secondary structures. Six rooted trees include the previously studied loop-stem (type 1) tree [25] and arc (type 6) tree [27], as well as four new tree representations (types 2, 3, 4 and 5). In particular, type 2 and type 4 trees can differentiate RNA secondary structures that are indistinguishable by other studied trees. We introduced tree-polynomials of RNA secondary structures by applying the tree distinguishing polynomial *P* [13] to type 1, 3, 5 and 6 trees and combining the newly developed tree distinguishing polynomial *Q* that better captures the loop group feature of RNA secondary structures with type 2 and 4 trees. These novel representations of RNA secondary structures enable fast and accurate analysis of extensive RNA secondary structures with modern data analytic tools. Moreover, they can be generalized for DNA nanotechnology and to analyze structures constructed by strand hybridization. In this paper, we first benchmark the performance of tree-polynomial representations in distinguishing different RNA secondary structures by an experiment of clustering non-coding RNA (ncRNA) secondary structures with distance-based clustering methods. The tree-polynomials accurately clustered ncRNA secondary structures with respect to the families of ncRNAs. We observed that ncRNA secondary structures in the same family but from different organisms have similar loop-stem relations and stem sizes but largely different loop sizes. This resulted in type 1 and type 3 tree-polynomials having the lowest mean misclassification rates and suggested that analyzing different structures requires different tree-polynomials. Next, we analyzed the link between secondary structures of newly transcribed RNA and R-loop formation. We found strong positive correlation between the probabilities of R-loop formation and the type 1, 2 and 4 scaled sums computed from corresponding tree-polynomials of the secondary structures of the nascent RNA. This suggests that we can use the scaled sums computed from the RNA sequence to predict R-loop formation. Furthermore, we identified the feature of an RNA secondary structure that contributes to a large coefficient sum. RNA secondary structures with large type 1, 2 and 4 coefficient sums share a common substructure consisting of many linear branches with short stems separated by multiple bulges and interior loops. We hypothesize that the short stems separated by bulges and interior loops in the linear branches make the base pairs in the branches easier to break, and base pair breakage aids the nascent RNA strand pairing with the DNA template strand and increases the probability of R-loop formation.

We used the co-transcriptional RNA folding model DrTransformer [1] to predict the secondary structures of 200-nucleotide long segments of the RNA transcript of the amplicon region of a plasmid. In doing this, we assumed that the secondary structure of an RNA segment does not affect that of other segments. Ideally, we would predict co-transcriptional RNA secondary structures of a full amplicon region. However, the amplicon regions of pFC8 and pFC53 are over 1000 nucleotides long. Performing the ideal experiment for each plasmid would require DrTransformer to process more than 500 RNA sequences with over 600 nucleotides. This is computationally expensive. DrTransformer suggests an input sequence shorter than 600 nucleotides. The computation time for a 200nt sequence is approximately 30 minutes on a single 3.2 GHz CPU core. Thus we restrict our sliding window to 200nt. The transcription process in this paper refers to the process of predicting the RNA secondary structures of the RNA segments *in silico* rather than a simulation of the *in vivo* transcription process of the amplicon region.

Finally, note that the tree-polynomials studied in this paper only work for RNA secondary structures without pseudoknots. In [21], the authors introduced an algebraic language for representing RNA secondary structures with pseudoknots. In order to study secondary structures with pseudoknots using polynomial-based methods, we can represent the secondary structures by directed acyclic graphs (DAGs). Recent studies have introduced complete polynomial invariants for some classes of DAGs in studying phylogenetic networks [10, 20, 31]. In future work we aim to modify the definitions of the polynomials and the rooted tree representations to capture more information about RNA secondary structures, for example, the order of stems and groups of unpaired nucleotides around a multiloop or the opening region. Furthermore, we can also apply the polynomials to other labeled, semi-labeled or unlabeled tree representations of RNA secondary structures. See [15] for an example of a complete polynomial invariant for labeled trees.

## Methods

### Background

#### RNA secondary structures

RNA molecules fold into secondary structures that can be described by paired and unpaired nucleotides. The paired nucleotides form double helices interspersed with structures of unpaired nucleotides [16]. We use the terms defined in [5] to describe different regions of an RNA secondary structure. See Figure 1 for an example of an RNA secondary structure and its regions and substructures.

A *stem* refers to an uninterrupted region of base pairs, and the two backbones are the *sides* of the stem. The space between two consecutive base pairs and their backbones is a *stem region*. A *hairpin loop* is the region bounded by a single strand of unpaired nucleotides and a stem, where both ends of the string of the unpaired nucleotides form a base pair at the stem. A *bulge* is the region bounded by a single strand of unpaired nucleotides and two stems, where both ends of the string of unpaired nucleotides attach to the same side of the two stems, and the other side of the two stems is an unbroken backbone. An *interior loop* is a region bounded by two strings of unpaired nucleotides and two stems, where two strings of unpaired nucleotides are flanked by the stems. A *multiloop* is the region bounded by *m* strings of unpaired nucleotides and *m* stems, for *m >* 2, such that the strings of unpaired nucleotides and the stems are alternately connected and form a cycle. The *opening region* is the unbounded region containing the 5’ and 3’ ends of the RNA molecule. A *loop* of an RNA secondary structure refers to a hairpin loop, a bulge, an interior loop, a multiloop or to the opening region. If a stem has nucleotides that belong to a loop, then we say the stem is *connected* to the loop. A stem connects two loops, one close to and the other away from the opening region. A *branch* is a substructure consisting of a stem and the loop away from the opening region together with all regions connected to the loop. A branch is *linear* if it contains only stems, bulges, interior loops and hairpin loops. A linear branch contains no multiloops.

#### Representations of RNA secondary structures

An RNA secondary structure is often encoded using a *dot-bracket notation* written in parallel with the RNA sequence. A dot represents an unpaired nucleotide, and a pair of brackets represent a pair of nucleotides. In Figure 1, we show the sequence and the dot-bracket notation of the secondary structure of a 41 nt long RNA molecule.

RNA secondary structures are also represented using graphs, especially rooted trees. An intuitive rooted tree representation is the *loop-stem tree*. Every vertex in a loop-stem tree represents a loop of the RNA secondary structure. The root vertex represents the opening region. An edge represents a stem of the RNA secondary structure. Two vertices *v*_1_ and *v*_2_ are connected by an edge *e* in the loop-stem tree if the two loops corresponding to *v*_1_ and *v*_2_ are connected by the stem represented by *e*. This and other graph representations are reviewed in [25].

The dot-bracket notation can be displayed as an arc diagram associated with a unique rooted tree [27], called the *arc tree* of the RNA secondary structure. Leaf vertices of an arc tree represent unpaired nucleotides in the secondary structure. An internal vertex represents either a loop or a stem region. Two internal vertices *v*_1_ and *v*_2_ are connected by an edge *e* in an arc tree if the loops or the stem regions represented by *v*_1_ and *v*_2_ are adjacent and separated by a single base pair. An edge *e* in an arc tree connects a leaf vertex *u* with an internal vertex *v* if the unpaired nucleotide represented by *u* is among the ones that bound the loop represented by *v*.

In Results, we systematically define six rooted tree representations of RNA secondary structures based on four structural features. These representations include the loop-stem tree and the arc tree.

It is possible for nucleotides in different loops to be connected via hydrogen bonds. The substructure constructed by such non-nested base pairs is called a *pseudoknot* [5]. See Figure 1 for an example. The bonds that define a pseudoknot are not reflected in the standard dot-bracket notation of an RNA secondary structure, the loop-stem tree, nor the arc tree. Other mathematical structures such as directed acyclic graphs are needed to represent pseudoknots. Here, we restrict our study to RNA secondary structures without pseudoknots.

#### A tree distinguishing polynomial

In [13], the author introduced a bivariate graph polynomial that distinguishes trees; we refer to it as *polynomial P*. A unique polynomial *P*(*T*, *x*, *y*) is associated with any given rooted tree *T* through the following process that recursively assigns bivariate polynomials to vertices of *T*, starting from the leaf vertices to the root vertex. Let *v* be a vertex in *T*. If *v* is a leaf vertex, then *P*(*v*, *x*, *y*) = *x*. If *v* is an internal vertex with *k* child vertices *u*_1_, *u*_2_, …, *u_k_*, then *P*(*v*, *x*, *y*) = *y* + Π*^k^ P*(*u_i_*, *x*, *y*). The polynomial for the entire tree *T* is the polynomial of the root vertex *r*, i.e. *P*(*T*, *x*, *y*) = *P*(*r*, *x*, *y*). Figure 2 illustrates the recursive process on type 1 and type 3 tree representations. Polynomial *P* can be directly applied to any rooted tree representation of RNA secondary structures without pseudoknots. In [13], the author also proved that polynomial *P* distinguishes trees, i.e. two trees *T*_1_ and *T*_2_ are isomorphic if and only if *P*(*T*_1_, *x*, *y*) = *P*(*T*_2_, *x*, *y*). Hence, polynomial *P* distinguishes the RNA secondary structures represented by the trees. Polynomial *P* is interpretable [13]. More precisely, every term in *P*(*T*, *x*, *y*) corresponds to a subtree of *T*, thus every term in *P*(*T*, *x*, *y*) describes a substructure of the RNA secondary structure represented by *T*.

#### Polynomial distance between trees

To compare and analyze RNA secondary structures represented by trees, we use a polynomial distance based on the Canberra distance as in [14]. A polynomial *P*(*T*, *x*, *y*) of degree *n* can be represented as an (*n* + 1) *×* (*n* + 1) coefficient matrix, where the entry *c*^(^*^i^*^,*j*)^ at the *j*-th row and the *i*-th column is the coefficient of the term *c*^(^*^i^*^,*j*)^ *x^i^y^j^* in *P*(*T*, *x*, *y*), for 0 *≤ i*, *j ≤ n*. Let *S* and *T* be two rooted trees and *b*^(^*^i^*^,*j*)^ and *c*^(^*^i^*^,*j*)^ be the coefficients of the corresponding terms in *P*(*S*, *x*, *y*) and *P*(*T*, *x*, *y*), respectively. The polynomial distance between *S* and *T* is defined by Formula (3), where the function *κ* is defined by Formula (2) in Results.

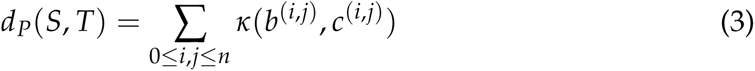

We call the distance defined above *polynomial P distance*. In Results, we generalize polynomial *P* and polynomial *P* distance to represent and compare newly introduced tree representations of RNA secondary structures.

### Data and experiments

#### Non-coding RNA secondary structures

We acquired data of RNA secondary structures from the bpRNA-1m database, which is constructed with the tool bpRNA that parses RNA structures from seven different sources [5]. We use one of the sources, Rfam, to construct a dataset of RNA secondary structures for our study. Rfam is a database that contains more than 25 families of non-coding RNAs (ncRNAs) [7]. Our dataset consists of 735 secondary structures without pseudoknots from seven families of ncRNAs chosen to have sequences of similar length with distinct secondary structures. We refer to this dataset as the *bpRNA-Rfam* dataset. See Supplementary Table 1-2.

Given a choice of tree-polynomial representation, we first cluster the 735 ncRNA secondary structures in the bpRNA-Rfam dataset with tree-polynomial distances, then we compare the clustering results with the true families of the ncRNA secondary structures. Specifically, we compute tree-polynomial distances between each pair of secondary structures in the bpRNA-Rfam dataset for a given tree-polynomial representation. We apply the k-medoids clustering algorithm [28] to the pairwise tree-polynomial distances. The k-medoids clustering algorithm is unsupervised, i.e. it assigns an integer between 1 and 7 to each ncRNA secondary structure. We use misclassification rates to compare the classes of RNA secondary structures determined by the clustering algorithm to the true families of the ncRNAs. The misclassification rate for an unsupervised clustering experiment can be computed by the majority rule [14]. Specifically, we associate a true family of ncRNA secondary structures with a cluster *i* if the majority of ncRNA secondary structures in cluster *i* are from the true family, where *i* is an integer between 1 and 7. Any secondary structure in cluster *i* that is not from the associated ncRNA family is misclassified. The misclassification rate of this clustering experiment is the percentage of misclassified secondary structures among the 735 ncRNA secondary structures in the bpRNA-Rfam dataset. Since the k-medoids algorithm is heuristic, we repeat the clustering experiment 1000 times for each tree-polynomial representation and compute the mean misclassification rate (MMR) over the 1000 experiments.

#### R-loop forming plasmids

We use sequence data of two double-stranded DNA plasmids that are known to form R-loops: pFC8 of length 3669 base pairs (bp) and pFC53 of length 3906 bp. These plasmids were studied in [29], and we only use the amplicon regions in our analysis. Specifically, the amplicon regions are between nucleotides 81-1512 in pFC8 and 81-1829 in pFC53. The *sequence of a plasmid* refers to the sequence of the amplicon region. We also use experimental data of the two circular plasmids with different topologies: negatively supercoiled and gyrase-treated (i.e. hyper-negatively supercoiled). These data were obtained by SMRF-seq based R-loop footprinting [17, 18] and reported in [29]. The circular plasmids were linearized after transcription and before SMRF-seq footprinting. Given a plasmid and its topology, the experimental data record the probability of R-loop formation at each nucleotide.

#### Sequence processing and structure prediction for R-loop forming plasmids

First, we convert the DNA sequence of a plasmid to its corresponding RNA sequence after transcription, which we refer to as the *RNA sequence of the plasmid*. Given an RNA sequence of a plasmid with *n* nucleotides, we take *m*-nucleotide long segments (*m < n*) starting at the 5’ end and shifting to the 3’ end in increments of one nucleotide. Specifically, for every integer *i* in the interval [1, *n − m* + 1], we take a segment between the *i*-th nucleotide and the (*i* + *m −* 1)-th nucleotide of the RNA sequence. For example, for pFC8, we have *n* = 1512 *−* 81 + 1 = 1432, and if we set *m* = 200, then the segments are 1-200 nt, 2-201 nt, 3-202 nt, … 1233-1432 nt of the RNA sequence. For each segment thus obtained, we use DrTransformer, an energy-based co-transcriptional RNA folding model [1], to predict the RNA secondary structures of the segment during transcription.

Given a segment of *m* nucleotides, DrTransformer produces a minimum-energy secondary structure at every transcription step. Specifically, each subsequence comprising the first to the *j*-th nucleotide of the segment corresponds to a minimum-energy secondary structure, for any integer *j* in the interval [1, *m*]. Thus, every segment has a series of *m* secondary structures with 1 to *m* nucleotides, which reflect how the secondary structure of the RNA segment changes during transcription. An RNA sequence of *n* nucleotides has *m*(*n − m* + 1) predicted secondary structures that can be located by two-dimensional coordinates (*i*, *j*), where *i* indicates the index of a segment in the RNA sequence and *j* indicates a transcription step. In this work, we set *m* = 200 to make it feasible to compute RNA secondary structures for thousands of RNA segments.

#### Computing tree-polynomial-based indices for R-loop forming plasmids

Given a plasmid’s *n*-nucleotide long RNA sequence, we compute the type *k* tree-polynomial representation for each of the *m*(*n − m* + 1) RNA secondary structures determined in the previous step, where *k* is an integer between 1 and 6. We compute the coefficient sum of the type *k* tree-polynomial that represents the secondary structure with coordinates (*i*, *j*) for every *i* and *j*. Thus, every secondary structure corresponds to a coefficient sum. We denote the coefficient sum of the secondary structure with coordinates (*i*, *j*) by *s*(*i*, *j*). To compare with the R-loop formation probabilities of a plasmid, we introduce the following variations of coefficient sums of secondary structures. For a transcription step *t*, there are *n − m* + 1 RNA secondary structures. The *normalized coefficient sum* or simply the *normalized sum* of the secondary structure at (*i*, *t*) is defined to be *s*(*i*, *t*)/ max_1_*_≤l≤n−m_*_+1_ *s*(*l*, *t*). Recall that the R-loop formation probability of a plasmid is a function of its sequence. For comparison, we define the *overlapping sum* of a plasmid at the transcription step *t* to imitate the experimental process of constructing the probability of R-loop formation [17, 18, 29]. The overlapping sum at the transcription step *t* is a function from the RNA sequence of the plasmid to the real numbers. We assign *s*(*i*, *t*) to every nucleotide between the *i*-th and the (*i* + *m −* 1)-th nucleotide (including the end nucleotides) for every integer *i* in the interval [1, *n − m* + 1]. Then, we add all coefficient sums assigned at every nucleotide of the RNA sequence. The result is the overlapping sum of the plasmid at transcription step *t*. See Supplementary Figure 8 for an illustration of the process. By dividing the overlapping sum of the RNA sequence at transcription step *t* by the maximum value of the overlapping sum, we have the *normalized overlapping sum* of the RNA sequence at transcription step *t*. By multiplying the normalized overlapping sum with the maximum R-loop formation probability of the plasmid, we have the *scaled overlapping sum* or simply the *scaled sum* of the RNA sequence of the plasmid. For a plasmid, we compare the scaled sum of the RNA sequence at transcription step *t* with the R-loop formation probability.

In summary, we construct sequence segments of the nascent RNA strand from the DNA sequences of the two plasmids, pFC8 and pFC53, and predict RNA secondary structures for the RNA segments. We compute the type *k* tree-polynomial representations of the RNA secondary structures for an integer *k* between 1 and 6. Thus, every secondary structure has a type *k* coefficient sum. In a transcription step *t*, every secondary structure has a type *k* normalized sum for comparing the relative difference to the secondary structure with the largest type *k* coefficient sum in the transcription step. For a transcription step *t*, we compute the type *k* overlapping sum, especially the type *k* scaled overlapping sum, from the type *k* coefficient sums of the secondary structures in the transcription step in order to compare with the probabilities of R-loop formation. Each plasmid has two probabilities of R-loop formation experimentally constructed with different DNA topologies: supercoiled and gyrase-treated. We compare the type *k* scaled sums with the two probabilities of R-loop formation. See Supplementary Figure 9 for a diagram displaying the entire computational pipeline.

#### Implementation and visualization

Code and data for experiments and analyses conducted in this paper are available at https://github.com/Arsuaga-Vazquez-Lab/RNA-Polynomial.

Interactive 3D figures for visualizing the normalized overlapping sums at every transcription step are available at https://pliumath.github.io/rloops.html.

## Acknowledgments

This work was supported by the National Science Foundation and the National Institutes of Health DMS/NIGMS award #2054347. The authors thank Ethan Holleman, Frédéric Chédin, Louxin Zhang and JianRong Yang for helpful discussion.

## Supplementary material

### Tree-polynomial representations

We study four important features of RNA secondary structures: loop-stem relation, loop size, stem size and loop group. All rooted tree representations record the loop-stem relation. There are eight possible ways to record combinations of the three other features. Six tree representations described in the main text. *Type 7* tree representations record loop-stem relation and loop group, and *type 8* tree representations record loop-stem relation, stem size and loop group. The first four tree representations are displayed in Figure 2 and the last four tree representations are displayed in Supplementary Figure 1.

Here, we describe how the eight tree representations are constructed from the RNA secondary structures. The loops in the RNA secondary structure are represented by the colored vertices in the tree representations in Figure 2 and Supplementary Figure 1. Vertex color corresponds to the type of loops; see Figure 1. The tree representations that record loop size (type 2, type 4, type 5 and type 6) have black leaf vertices that represent unpaired nucleotides around loops. Let *L* be a loop and *n*_1_, *n*_2_, …, *n_k_*be the unpaired nucleotide around the loop *L*. The black leaf vertices that represent *n*_1_, *n*_2_, …, *n_k_* are descendants of the colored vertex that represents the loop *L*. The tree representations that record stem size (type 3, type 4, type 6 and type 8) have gray internal vertices that represent stem regions. A stem with *n* base pairs in the RNA secondary structure is represented by a path only consisting of *n −* 1 gray vertices in the tree representations. Since stems do not have unpaired nucleotides, the gray vertices in the tree representations do not have child vertices that are leaf vertices in the trees. The tree representations that record both loop size and loop group (type 2, and type 4) have black square artificial vertices representing the groups of unpaired nucleotides around each loop. Suppose that *g* is a group of the unpaired nucleotides *n*_1_, *n*_2_, …, *n_k_* around a loop *L*. The black square artificial vertex that represents group *g* is a child vertex of the colored vertex representing *L*. The black leaf vertices that represent unpaired nucleotides *n*_1_, *n*_2_, …, *n_k_* are child vertices of the black square artificial vertex that represents the group *g*. In Supplementary Figure 2, we show an example of an RNA secondary structure that can be distinguished by the type 2 and the type 4 tree representations but not the other six tree representations. The two distinct 52-nucleotide long RNA secondary structures have the same loop-stem relation, the corresponding stems have the same size and the corresponding loops have the same number of unpaired nucleotides. The only difference between the secondary structures is how the unpaired nucleotides are grouped around each loop.

The tree representations that record loop group but not loop size (type 7 and type 8) have the artificial vertices that represent groups of unpaired nucleotides as leaf vertices, so we represent them with the black round vertices; see Supplementary Figure 1. Type 7 and the type 8 tree representations provide little additional information about the RNA secondary structure compared to the type 1 and type 3 tree representations respectively. This is because the groups of unpaired nucleotides are divided by branches around a loop. Type 7 and type 8 tree representations have the following properties: a hairpin loop must only have one artificial vertex as a child and leaf vertex; a bulge must have two child vertices including one and only one artificial vertex as a leaf vertex; an interior loop must have three child vertices including two artificial vertices as leaf vertices; a multiloop with *n* branches must have 2*n –* 1 child vertices with *n* artificial vertices as leaf vertices. The only additional information that a type 7 or a type 8 tree representation can provide when compared to a type 1 or a type 3 tree representation is whether the nucleotides at the 5’ and the 3’ ends are paired or unpaired. Suppose that the opening region of an RNA secondary structure has *n* branches. If both ends are paired, then the root vertex has *n −* 1 artificial vertices as child and leaf vertices. If one end is paired, then the root vertex has *n* artificial vertices as child and leaf vertices. If both ends are unpaired, then the root vertex has *n* + 1 artificial vertices as child and leaf vertices. Moreover, the type 7 and type 8 tree-polynomial representations of an RNA secondary structure have the same coefficient sums as the type 5 and type 6 tree-polynomial representations, respectively. This is because leaf vertices contribute only to the exponents and not to the coefficients. It is for these reasons that we do not discuss type 7 and type 8 tree-polynomial representations in the main text. Type 7 and type 8 tree-polynomial representations also perform the worst in clustering the non-coding RNA secondary structures in the bpRNA-Rfam dataset, with mean misclassification rates 24.32% and 22.16% respectively.

**Supplementary Figure 1:**
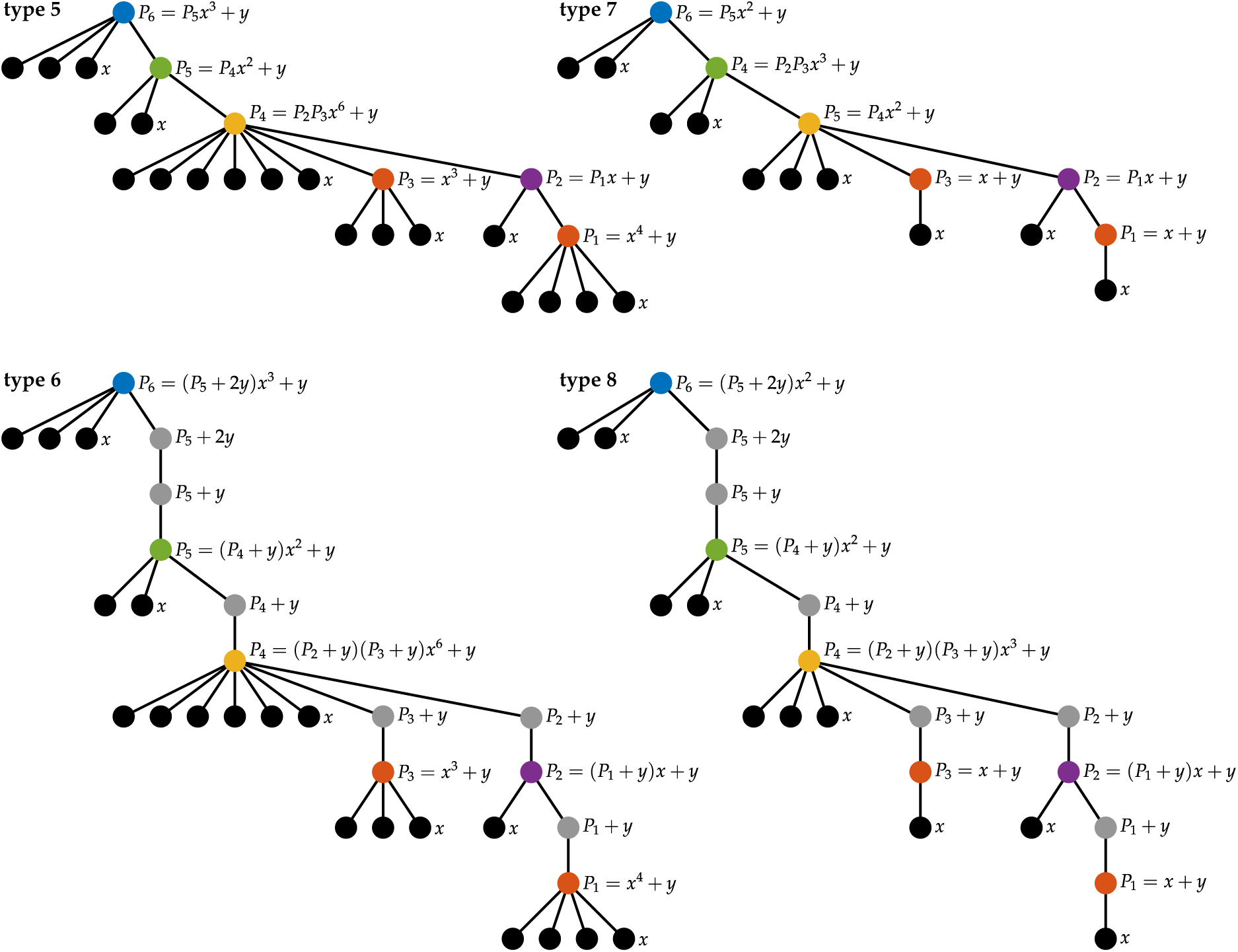
Rooted tree and polynomial representations of RNA secondary structures. The figure shows the last four rooted tree representations and their corresponding polynomial representations of the RNA secondary structure displayed in Figure 1. Vertices in the trees are colored based on the loops or stem regions that they represent, and the black round vertices represent unpaired nucleotides. The leaf vertices in type 7 and type 8 tree representations represent artificial vertices introduced for grouping unpaired nucleotides. Alongside every tree representation, the recursive process of computing the corresponding polynomial from the leaf vertices to the root vertex is displayed. The polynomial at the root vertex of a rooted tree is the polynomial that represents the tree.

**Supplementary Figure 2:**
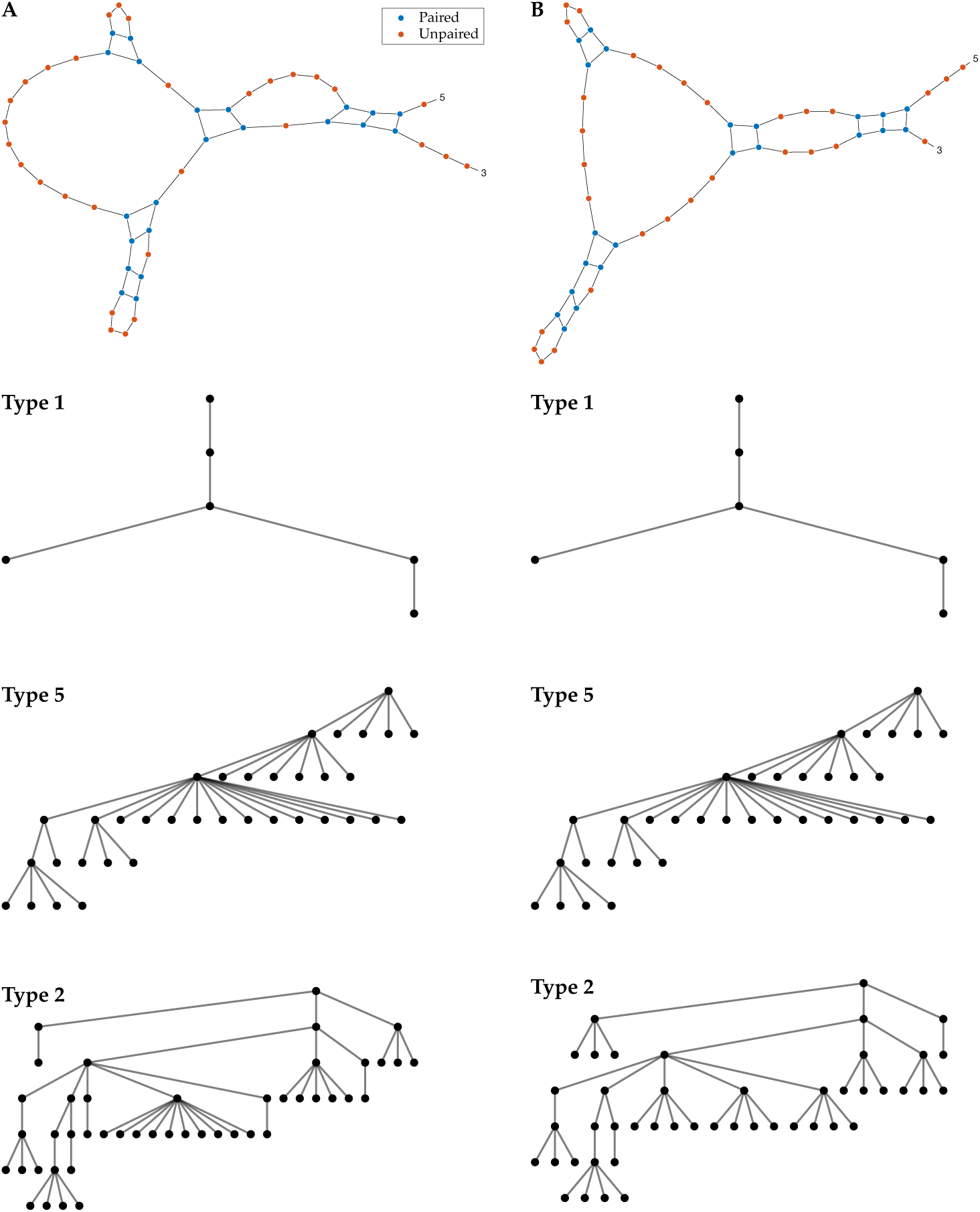
An example of a pair of RNA secondary structures that requires type 2 and type 4 tree representations to distinguish. Panel A and panel B show a pair of 52-nucleotide long RNA secondary structures that are the same in loop-stem relation, loop size and stem size but different in loop group. Each panel also displays the corresponding type 1, type 5 and type 2 tree representations, where only type 2 tree representations distinguish between the RNA secondary structures.

We use polynomial *Q* to define type 2 and type 4 tree-polynomial representations. Other types of tree representations use the previously studied tree distinguishing polynomial *P*. The recursive processes of computing the polynomials from the tree representations are illustrated in Figure 2 and Supplementary Figure 1. Given a rooted tree *T*, note that if we take *z* = *y* in a polynomial *Q*(*T*, *x*, *y*, *z*), then we obtain the polynomial *P*(*T*, *x*, *y*). Since the polynomial *P* distinguishes trees, the polynomial *Q* also distinguishes trees. Mathematically, two trees are isomorphic if and only if they have the same polynomial *Q*.

In Supplementary Figure 3, we show the average computational time needed to compute each tree-polynomial representation for a secondary structure in the bpRNA-Rfam dataset. The average length of an ncRNA secondary structure in the dataset is 171.97 nucleotides; see Supplementary Table 1. As expected, computing the multivariate polynomial *Q* is more expensive than computing the bivariate polynomial *P*.

### Clustering RNA secondary structures

Basic information of the seven families of non-coding RNA (ncRNA) secondary structures in the bpRNA-Rfam dataset is listed in Supplementary Table 1. More detailed information about an individual ncRNA secondary structure in the dataset can be found at the bpRNA-1m database (https://bprna.cgrb.oregonstate.edu), where each RNA secondary structure is associated with a bpRNA ID. Supplementary Table 2 lists the bpRNA IDs of the ncRNA secondary structures in the bpRNA-Rfam dataset.

In Supplementary Figure 4-5, we visualize pairwise polynomial distances between the tree-polynomial representations of the ncRNA secondary structures with multidimensional scaling (MDS).

In Supplementary Figure 6-7, we display examples of ncRNA secondary structures from the 5.8S ribosomal RNA family and the U12 minor spliceosomal RNA family in the bpRNA-Rfam dataset. Examples of other secondary structures in the bpRNA-Rfam dataset can be found at the bpRNA-1m database with their bpRNA ID. The We observe that ncRNA secondary structures in the same family from different organisms can display large differences in loop size, while their loop-stem relation and stem size are consistent. This partially explains why type 3 and type 1 tree representations perform the best in clustering the ncRNA secondary structures of the bpRNA-Rfam dataset.

### RNA secondary structures and R-loop formation

In Methods, we describe the process of computing the overlapping sum, the normalized overlapping sum and the scaled overlapping sum based on the coefficient sums of type *k* of tree-polynomial representations of RNA segments for an integer *k* between 1 and 8. This process is visualized in Supplementary Figure 8 with a toy example. In this toy example, the RNA sequence is of *n* = 14 nucleotides, and the length of a segment is *m* = 5 nucleotides. So, we have 10 segments for the RNA sequence in increments of one nucleotide. The segments are depicted in panel A. For every segment, DrTransformer predicts *m* = 5 co-transcriptional RNA secondary structures. The secondary structures have length from one nucleotide to *m* = 5 nucleotides. The predicted co-transcriptional RNA secondary structures for the first and the last segment are displayed in panel A. The parameters used in DrTrasnformer are all at default values in this paper. We take the secondary structures for all segments at transcription step *j* = 5. For the secondary structure of the *i*-th segment, we compute its tree-polynomial representation and the coefficient sum *s*(*i*, *j*). We put the number *s*(*i*, *j*) at every nucleotide in the *i*-th segment (panel B). The overlapping sum of the RNA sequence is computed by vertically adding all numbers at each nucleotide (panel B). We divide the overlapping sum by its maximum value (29 in the example) to obtain the normalized overlapping sum of the RNA sequence (the yellow curve in panel B). Then, we multiply the normalized overlapping sum with the highest probability of R-loop formation to obtain the scaled overlapping sum of the RNA sequence (the black curve in panel B).

**Supplementary Figure 3:**
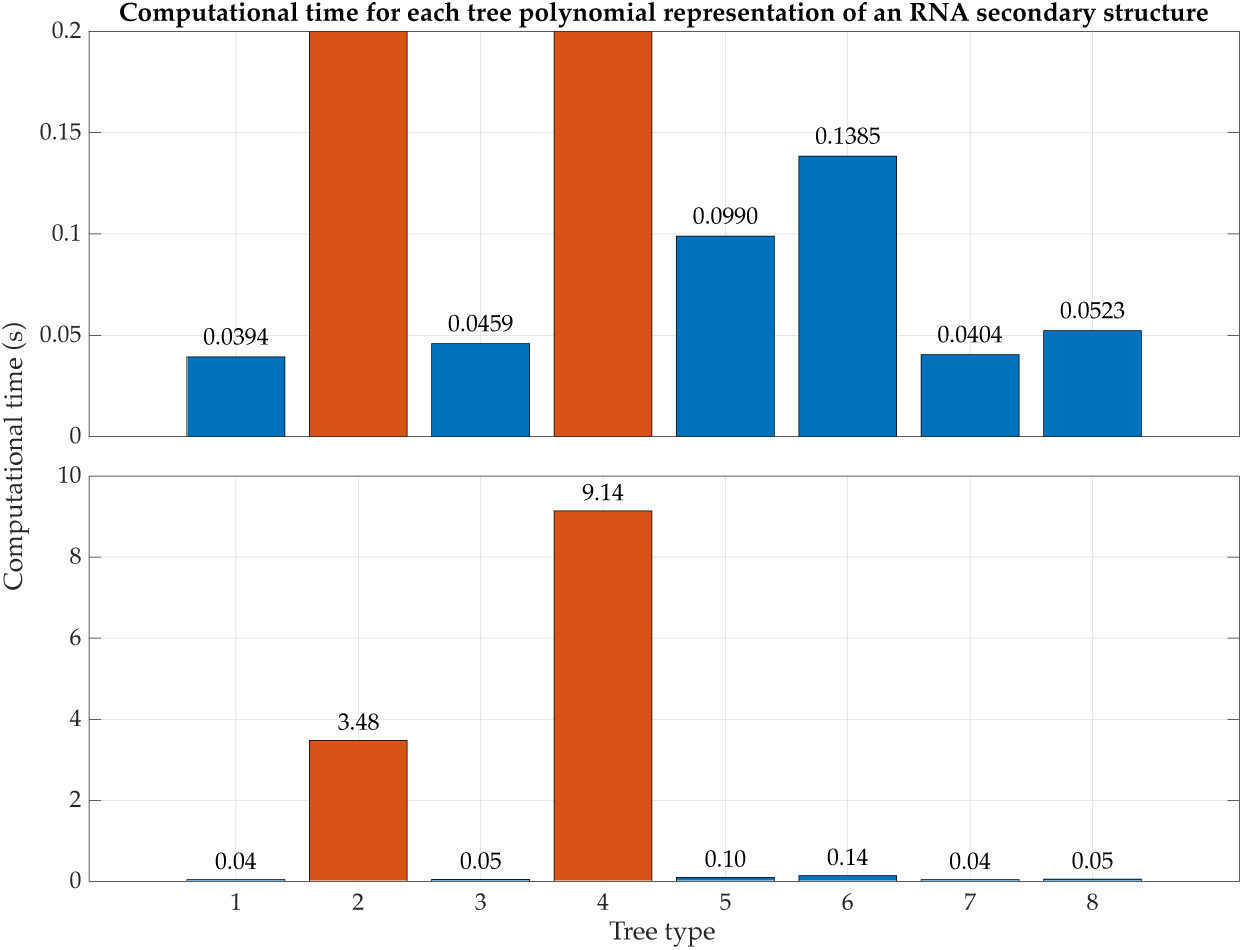
Average computational time of each tree-polynomial representation for an RNA secondary structure. Each bar shows the average time spent computing the corresponding tree-polynomial representations of the 735 ncRNA secondary structures (with average length 171.97 nt) in the bpRNA-Rfam dataset on a single 3.2 GHz CPU core. Blue bars indicate that the representations use polynomial *P*, and red bars indicate that the representations use polynomial *Q*. The top panel shows a more detailed comparison of the blue bars.

**Supplementary Table 1:**
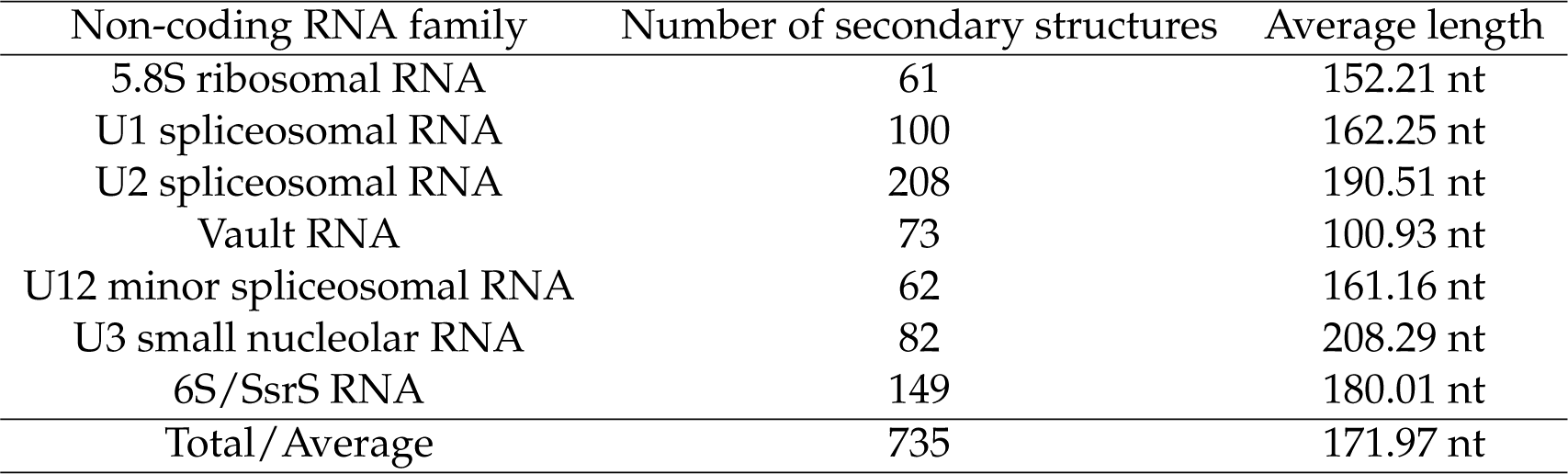
The bpRNA-Rfam dataset of non-coding RNA (ncRNA) secondary structures. The first column shows the seven Rfam ncRNA families selected in the study. The second column shows the number of RNA secondary structures without pseudoknots in each family. The third column shows the average length of the ncRNA sequences in each family.

**Supplementary Table 2:** The bpRNA IDs of ncRNA secondary structures in the bpRNA-Rfam dataset. The table shows the bpRNA ID of the secondary structures from the seven ncRNA families in the Rfam dataset. The asterisk superscript indicates that there are five U3 small nucleolar RNA secondary structures (with ID 2946, 2951, 2965, 2970 and 2998) that have pseudoknots and are not included in the bpRNA-Rfam dataset.

| Non-coding RNA family | bpRNA ID |
| --- | --- |
| 5.8S ribosomal RNA | bpRNA_RFAM_713 to bpRNA_RFAM_773 |
| U1 spliceosomal RNA | bpRNA_RFAM_774 to bpRNA_RFAM_873 |
| U2 spliceosomal RNA | bpRNA_RFAM_874 to bpRNA_RFAM_1801 |
| Vault RNA | bpRNA_RFAM_2036 to bpRNA_RFAM_2108 |
| U12 minor spliceosomal RNA | bpRNA_RFAM_2109 to bpRNA_RFAM_2170 |
| U3 small nucleolar RNA | bpRNA_RFAM_2941 to bpRNA_RFAM_3027* |
| 6S/SsrS RNA | bpRNA_RFAM_3028 to bpRNA_RFAM_3176 |

**Supplementary Figure 4:**
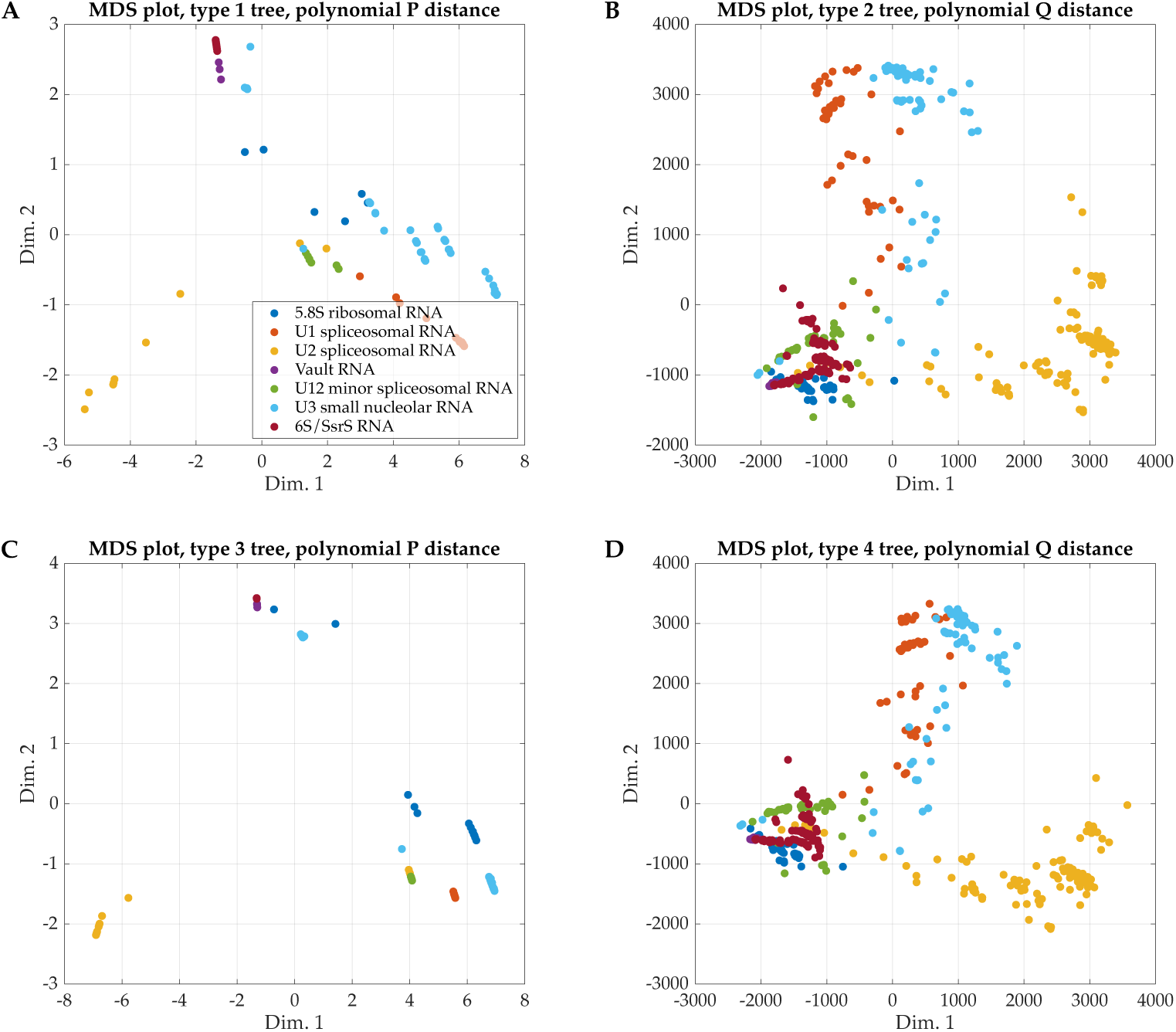
Visualization of pairwise tree-polynomial distances between non-coding RNA (ncRNA) secondary structures in the bpRNA-Rfam dataset. The top two panels show the MDS plots of the pairwise polynomial *P* distances between type 1 (panel A) and type 2 (panel B) tree-polynomial representations of the 735 ncRNA secondary structures in the bpRNA-Rfam dataset. The bottom panels show the analogous MDS plots of the pairwise polynomial *Q* distances between type 3 (panel C) and type 4 (panel D) tree-polynomial representations. Each dot in a panel represents an ncRNA secondary structure of the ncRNA family corresponding to its color. We observe distinct clusters for each of the seven families of ncRNAs in all four plots. The plot for type 3 tree-polynomial representations corresponds to the best clustering results, with only two U3 small nucleolar RNAs (cyan) close to other clusters.

**Supplementary Figure 5:**
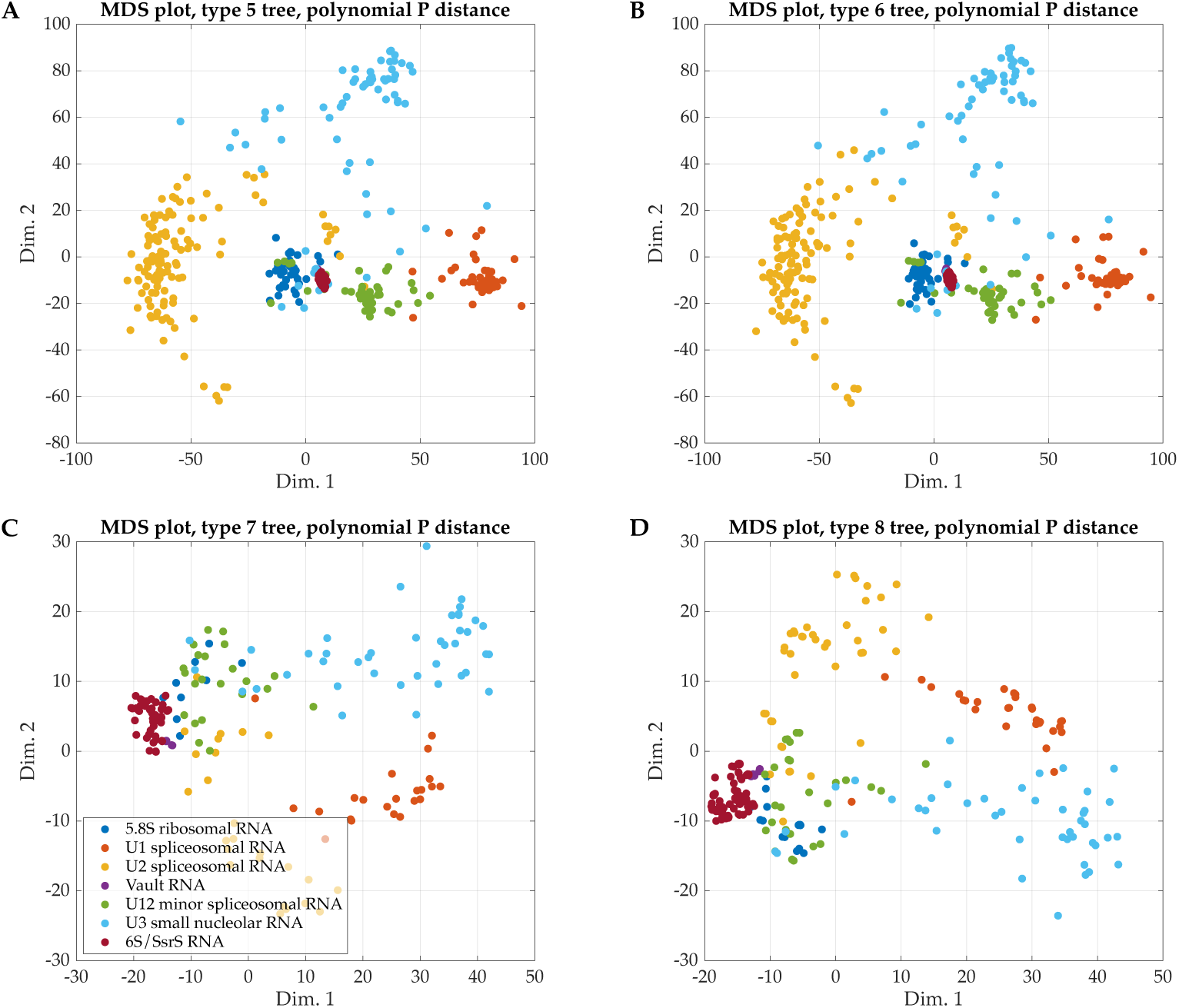
Visualization of pairwise tree-polynomial distances between non-coding RNA (ncRNA) secondary structures in the bpRNA-Rfam dataset. The figure shows the MDS plots of the pairwise polynomial *P* distances between type 5 (panel A) and type 6 (panel B) tree-polynomial representations of the 735 ncRNA secondary structures in the bpRNA-Rfam dataset, and the MDS plots of the pairwise polynomial *P* distances between type 7 (panel C) and type 8 (panel D) tree-polynomial representations of the ncRNA secondary structures in the bpRNA-Rfam dataset. Each dot in a panel represents an ncRNA secondary structure of the ncRNA family corresponding to its color.

**Supplementary Figure 6:**
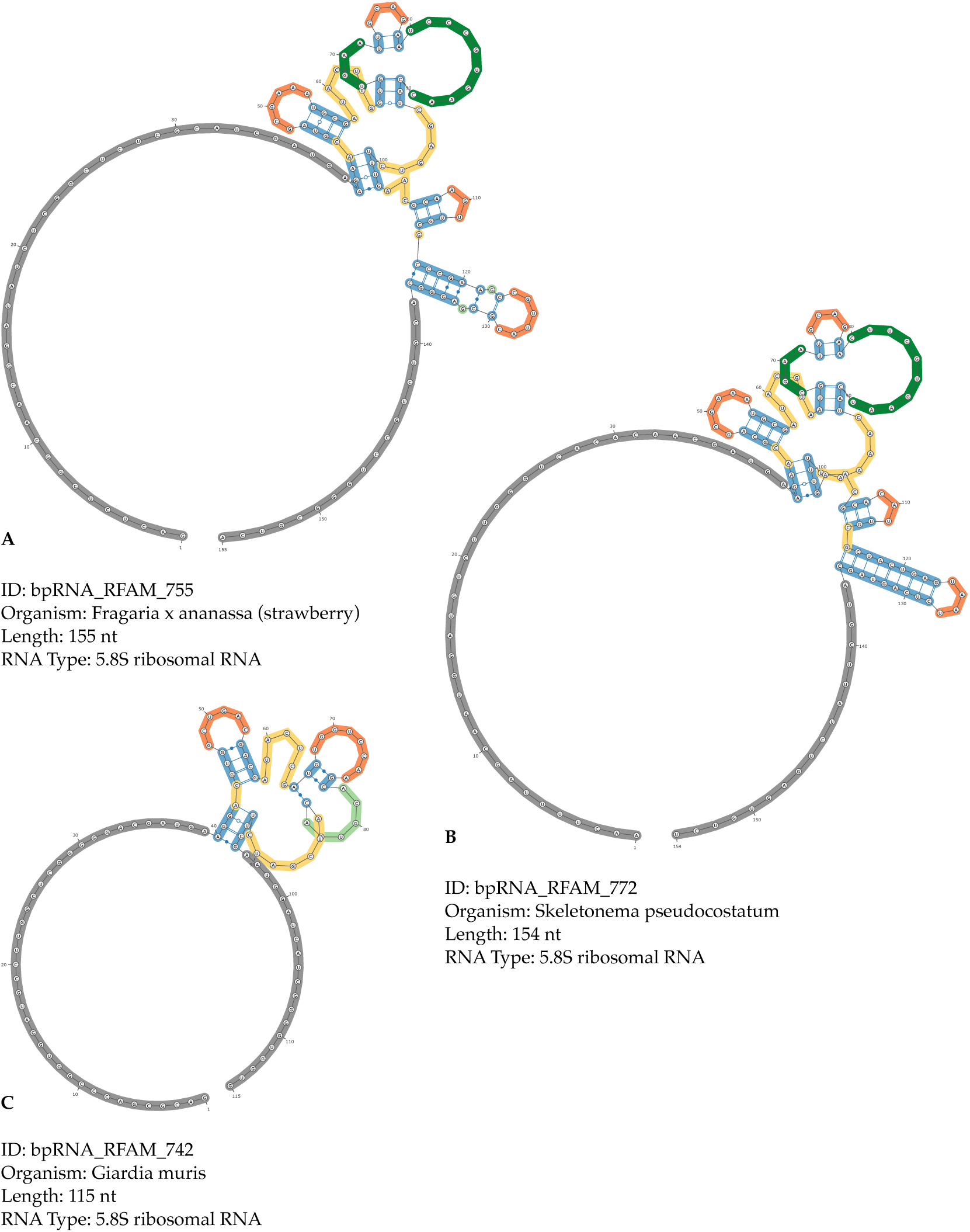
Examples of secondary structures from the same ncRNA family in the bpRNA-Rfam dataset. This figure shows examples of 5.8S ribosomal RNA secondary structures in the bpRNA-Rfam dataset. We obtained the secondary structure diagrams and information from bpRNA-1m.

**Supplementary Figure 7:**
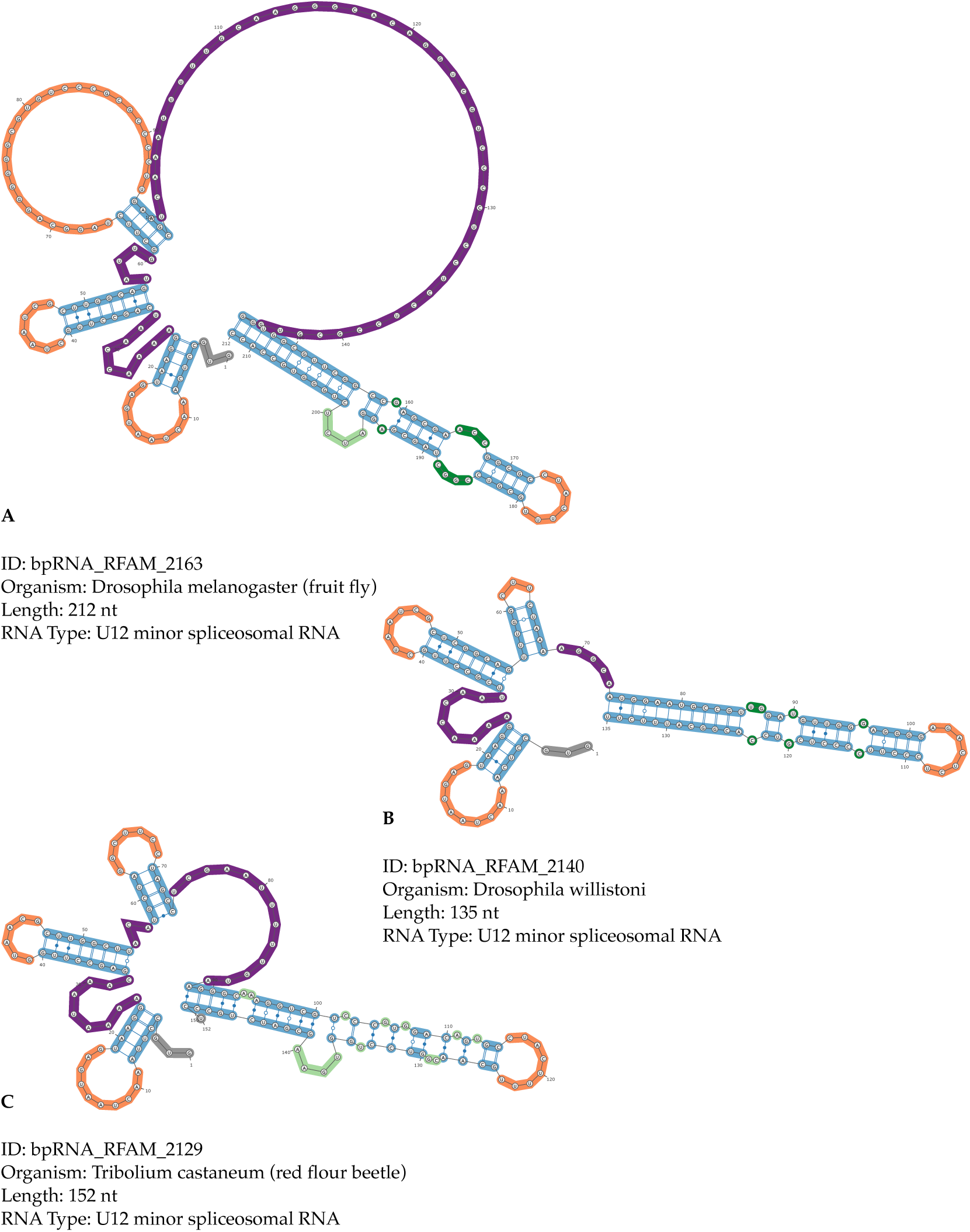
Examples of secondary structures from the same ncRNA family in the bpRNA-Rfam dataset. This figure shows examples of U12 minor spliceosomal RNA secondary structures in the bpRNA-Rfam dataset. We obtained the secondary structure diagrams and information from bpRNA-1m.

In Methods, we also describe pipeline of the computational experiment for analyzing the link between the secondary structures of the nascent RNA strand and the probabilities of R-loop formation. In Supplementary Figure 9, we visualize the computational pipeline of how we use the data of the two plasmids to conduct the experiment.

In Supplementary Figure 10, we replace the probabilities of R-loop formation of supercoiled plasmids in Figure 4 with the probabilities of R-loop formation of gyrase-treated plasmids, and show the correlations with the type 1 and type 4 scaled sums. The R-loop formation probabilities of the gyrase-treated plasmids have two outstanding peaks, while the R-loop formation probabilities of the supercoiled plasmids have only one outstanding peaks. The higher peaks indicate the positions of the major R-loop forming sequence cluster and the lower peaks indicate the positions of the minor R-loop forming sequence cluster of the two plasmids.

In Supplementary Figure 11, we show the correlations between probabilities of R-loop formation of supercoiled and gyrase-treated plasmids and the type 2 scaled sums. The type 2 scaled sums are similar to the type 4 scaled sums for the two plasmids. The difference between the two tree representations is that type 4 tree representations record stem size while the type 2 ones do not. In Supplementary Figure 12, we show the secondary structures of RNA segments that have the two largest type 2 coefficient sums and the corresponding type 2 tree representations. In Supplementary Figure 13, we show the secondary structures of RNA segments that have the two largest type 4 coefficient sums and the corresponding type 4 tree representations.

In Supplementary Figure 14, we show the correlations between R-loop formation probabilities of supercoiled and gyrase-treated plasmids and the type 3 scaled sums. Note that the type 3 tree-polynomial representation records the stem size but not the loop size or loop group with the bivariate polynomial *P*. The type 3 scaled sums are not strongly correlated with the probabilities of R-loop formation. The type 3 tree representation of a linear branch is a path with vertices representing bubbles and stem regions, which are not differentiated by the tree-polynomial representations. This mixing of information is partially why the type 3 scaled sums and the R-loop formation probabilities are not strongly correlated. The RNA secondary structures with the two largest type 3 coefficient sums are displayed in Supplementary Figure 15-16, where both the type 1 and the type 3 tree representations are shown. We observe that most of the linear branches in these secondary structures do not have many bubbles, and the large type 3 coefficient sums are contributed by the vertices that represent stem regions.

In Supplementary Figure 17, we show the secondary structures of RNA segments that have the two largest type 1 coefficient sums near the 3’ end of the amplicon region, where few R-loops form. The corresponding type 1 tree representations are displayed following the secondary structures. In Supplementary Figure 17, we show the secondary structures of RNA segments that have the two largest type 1 coefficient sums in the minor R-loop forming cluster and the corresponding type 1 tree representations are displayed following the secondary structures.

The results discussed so far only cover the scaled sums that have the highest Pearson’s correlation coefficients (PCCs) with the probabilities of R-loop formation in the last 10 transcription steps. Analogous data of the PCCs for type 5 to type 8 tree-polynomial representations are listed in Supplementary Table 3. As previously discussed, the type 7 and the type 8 tree-polynomial representations of an RNA secondary structure have the same coefficient sums as the type 5 and type 6 tree-polynomial representations of the secondary structure respectively. Hence, the type 7 and the type 8 scaled sums of the plasmids are identical to the type 5 and the type 6 scaled sums of the plasmids, and they have the same PCCs. In Supplementary Table 4, we list the PCCs of the scaled sums that are most correlated with the probabilities of R-loop formation in all transcription steps. The interactive 3D figures available online at https://pliumath.github.io/rloops.html show how the normalized overlapping sums change with respect to the transcriptional process of the RNA segments. We also observe in the interactive 3D figures that the values of normalized overlapping sums become relatively stable and the strong correlations appear at a very early stage of the transcriptional process, when the secondary structures of the segments have fewer than 50 nucleotides. The difference between the secondary structures with large coefficient sums and the ones with small coefficient sums are also captured by the tree-polynomial representations of the fewer-than-50-nucleotide long secondary structures. This suggests that the fewer-than-50-nucleotide long secondary structures already possess the feature for large coefficient sums, which potentially promotes R-loop formation.

**Supplementary Figure 8:**
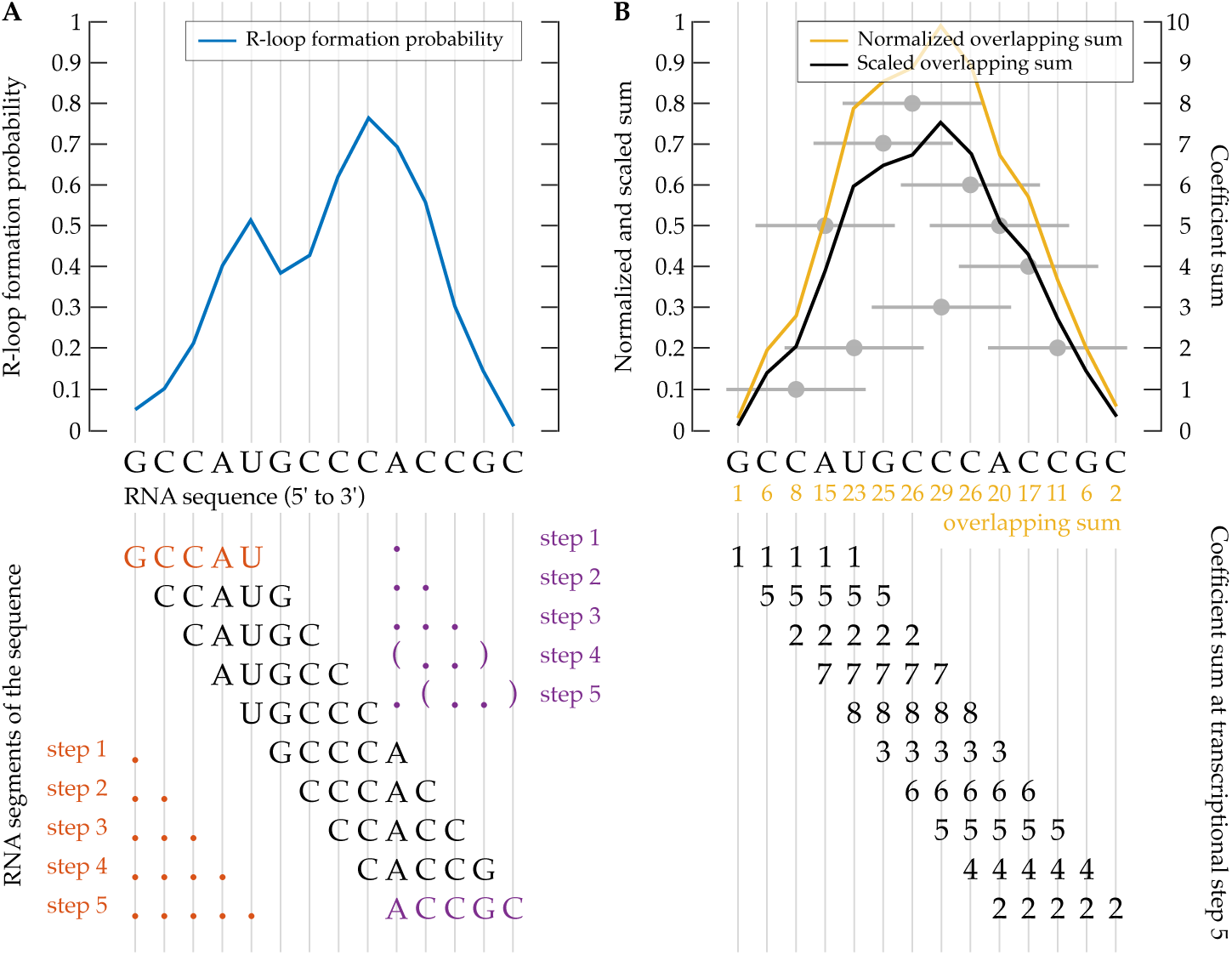
Visualization of the process for computing normalized and scaled overlapping sums. Panel A shows the probability of R-loop formation (blue) as a function of the RNA sequence, segments of the RNA sequence and the lists of RNA secondary structures (red and purple) of the first and the last RNA segments. Panel B shows the coefficient sums (black numbers) and the normalized sums (gray dots and horizontal lines) of the RNA segments at a transcription step and the process of computing the overlapping sum (yellow numbers below the sequence), normalized overlapping sum (yellow curve) and the scaled sum (black curve) of the RNA sequence from the coefficient sums.

**Supplementary Table 3:** The highest Pearson’s correlation coefficients between the scaled sums and the probabilities of R-loop formation of the last 10 transcription steps. The first columns show the types of tree-polynomial representations. The last four columns show the highest Pearson’s correlation coefficients (PCCs) between the scaled sums of the tree-polynomial representations and the probabilities of R-loop formation of the corresponding plasmids in the last 10 transcription steps. Inside the parentheses after the PCCs are the transcription steps of the corresponding scaled sums.

| Type | pFC8 supercoiled | pFC8 gyrase | pFC53 supercoiled | pFC53 gyrase |
| --- | --- | --- | --- | --- |
| 1 | 0.77 (step 199) | 0.62 (step 199) | 0.79 (step 196) | 0.81 (step 196) |
| 2 | 0.96 (step 199) | 0.77 (step 200) | 0.96 (step 192) | 0.88 (step 192) |
| 3 | -0.30 (step 198) | -0.28 (step 198) | -0.19 (step 200) | -0.11 (step 200) |
| 4 | 0.97 (step 199) | 0.78 (step 200) | 0.97 (step 195) | 0.89 (step 191) |
| 5 | 0.60 (step 197) | 0.48 (step 197) | 0.64 (step 195) | 0.66 (step 199) |
| 6 | -0.31 (step 198) | -0.34 (step 198) | -0.21 (step 200) | -0.18 (step 194) |
| 7 | 0.60 (step 197) | 0.48 (step 197) | 0.64 (step 195) | 0.66 (step 199) |
| 8 | -0.31 (step 198) | -0.34 (step 198) | -0.21 (step 200) | -0.18 (step 194) |

**Supplementary Figure 9:**
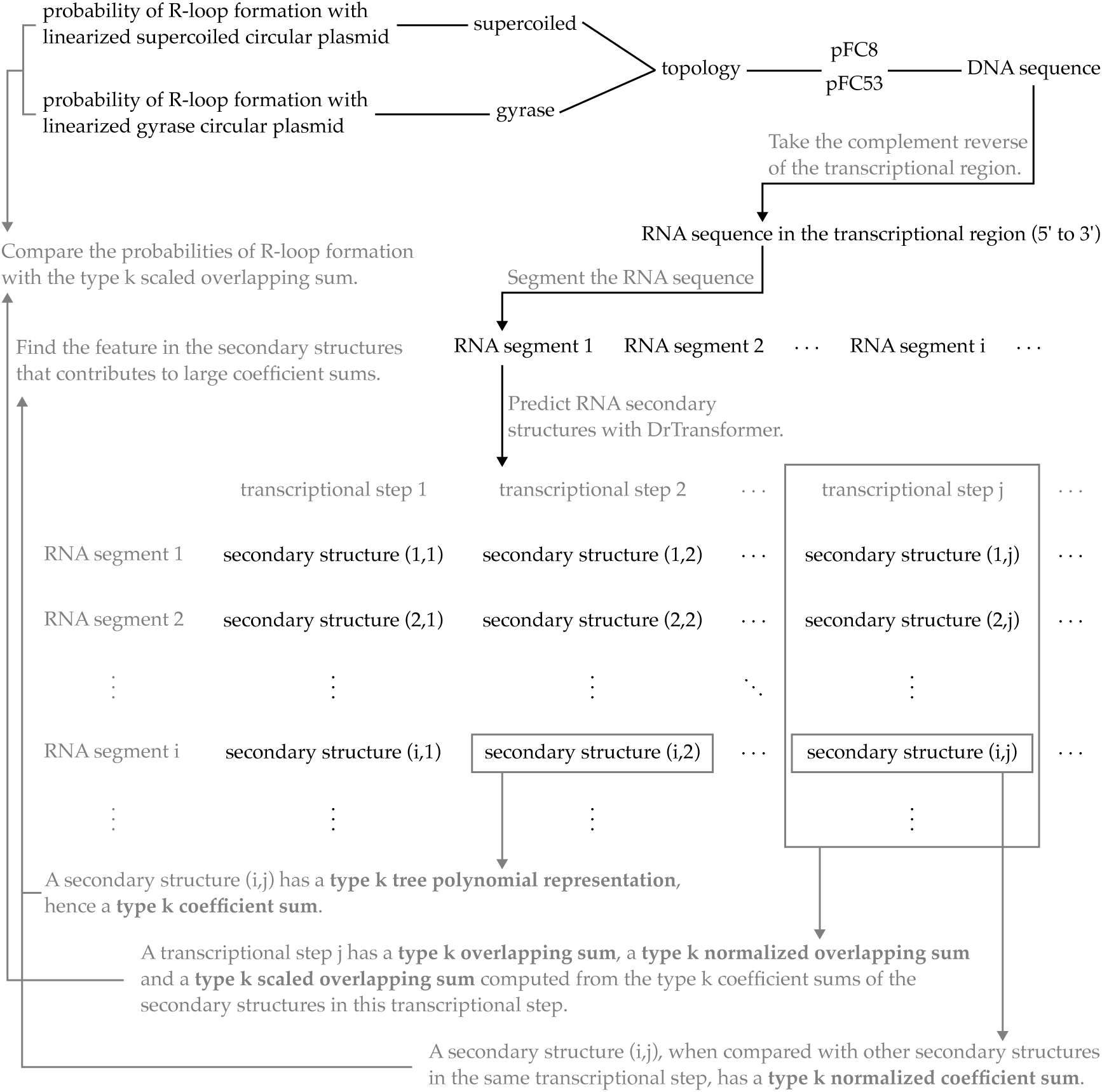
A diagram of the computational pipeline for analyzing the link between RNA secondary structures and R-loop formation. This diagram shows how we process the DNA sequence data of the two plasmids and compare the predicted RNA secondary structure with the probabilities of R-loop formation.

**Supplementary Table 4:** The highest Pearson’s correlation coefficients between the scaled sums and the probabilities of R-loop formation in all transcription steps. The first columns show the types of tree-polynomial representations. The last four columns show the highest Pearson’s correlation coefficients (PCCs) between the scaled sums of the tree-polynomial representations and the probabilities of R-loop formation of the corresponding plasmids over all transcription steps. Inside the parentheses after the PCCs are the transcription steps of the corresponding scaled sums.

| Type | pFC8 supercoiled | pFC8 gyrase | pFC53 supercoiled | pFC53 gyrase |
| --- | --- | --- | --- | --- |
| 1 | 0.77 (step 199) | 0.62 (step 181) | 0.79 (step 196) | 0.81 (step 196) |
| 2 | 0.96 (step 199) | 0.77 (step 200) | 0.97 (step 134) | 0.90 (step 121) |
| 3 | 0.32 (step 20) | 0.42 (step 16) | -0.21 (step 167) | 0.27 (step 9) |
| 4 | 0.97 (step 199) | 0.78 (step 200) | 0.97 (step 150) | 0.90 (step 153) |
| 5 | 0.62 (step 168) | 0.49 (step 181) | 0.71 (step 147) | 0.72 (step 156) |
| 6 | 0.32 (step 20) | 0.39 (step 16) | 0.24 (step 67) | 0.27 (step 9) |
| 7 | 0.62 (step 168) | 0.49 (step 181) | 0.71 (step 147) | 0.72 (step 156) |
| 8 | 0.32 (step 20) | 0.39 (step 16) | 0.24 (step 67) | 0.27 (step 9) |

**Supplementary Figure 10:**
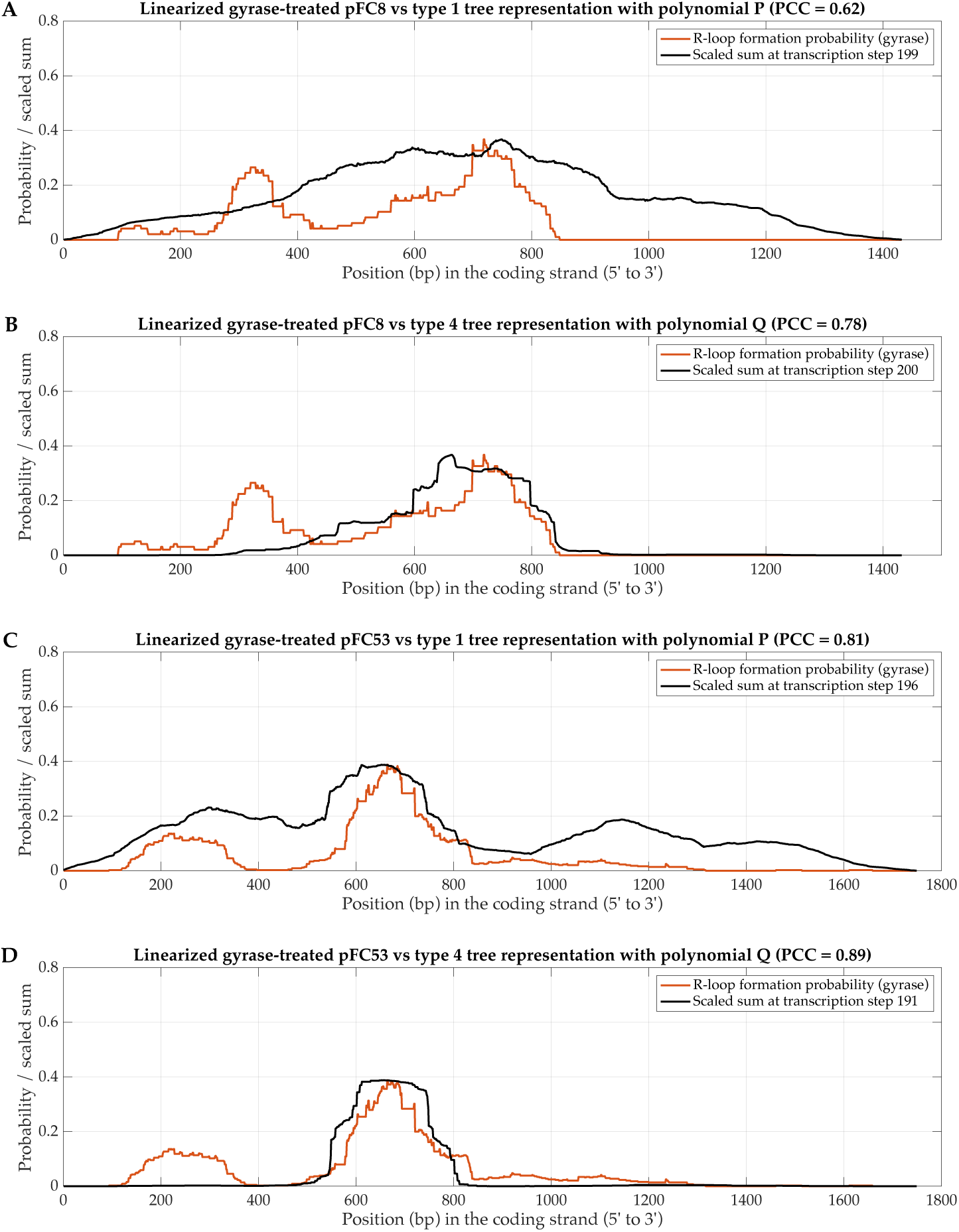
The correlations between the scaled sums and the R-loop formation probabilities of linearized gyrase-treated pFC8 and pFC53 plasmids. The figure shows the experimental probability of R-loop formation for the linearized gyrase-treated pFC8 plasmid with the type 1 (panel A) and the type 4 (panel B) scaled sums, and the experimental probability of R-loop formation for the linearized gyrase-treated pFC53 plasmid with the type 1 (panel C) and the type 4 (panel D) scaled sums. The experimental probabilities of R-loop formation are from [29]. The displayed scaled sums have the highest PCC in the last 10 transcription steps.

**Supplementary Figure 11:**
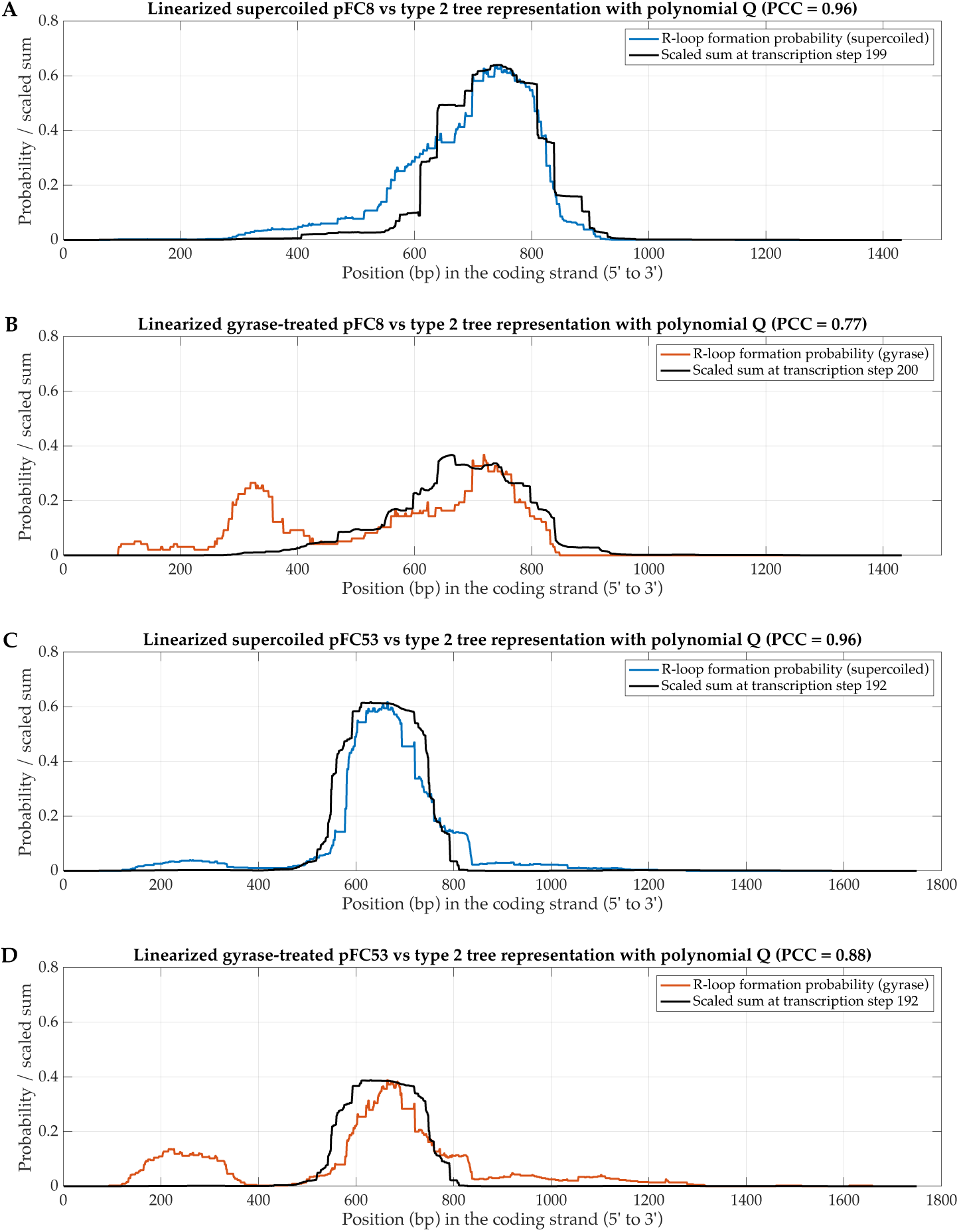
The correlations between the type 2 scaled sums and the R-loop formation probability of pFC8 and pFC53 plasmids. The figure shows the correlations between the type 2 scaled sums (with the highest PCC in the last 10 transcription steps) and the R-loop formation probabilities of the linearized supercoiled pFC8 plasmid (panel A) and of the linearized gyrase-treated pFC8 plasmid (panel B), and the correlations between the type 2 scaled sums (with the highest PCC in the last 10 transcription steps) and the R-loop formation probabilities of the linearized supercoiled pFC53 plasmid (panel C) and of the linearized gyrase-treated pFC53 plasmid (panel D).

**Supplementary Figure 12:**
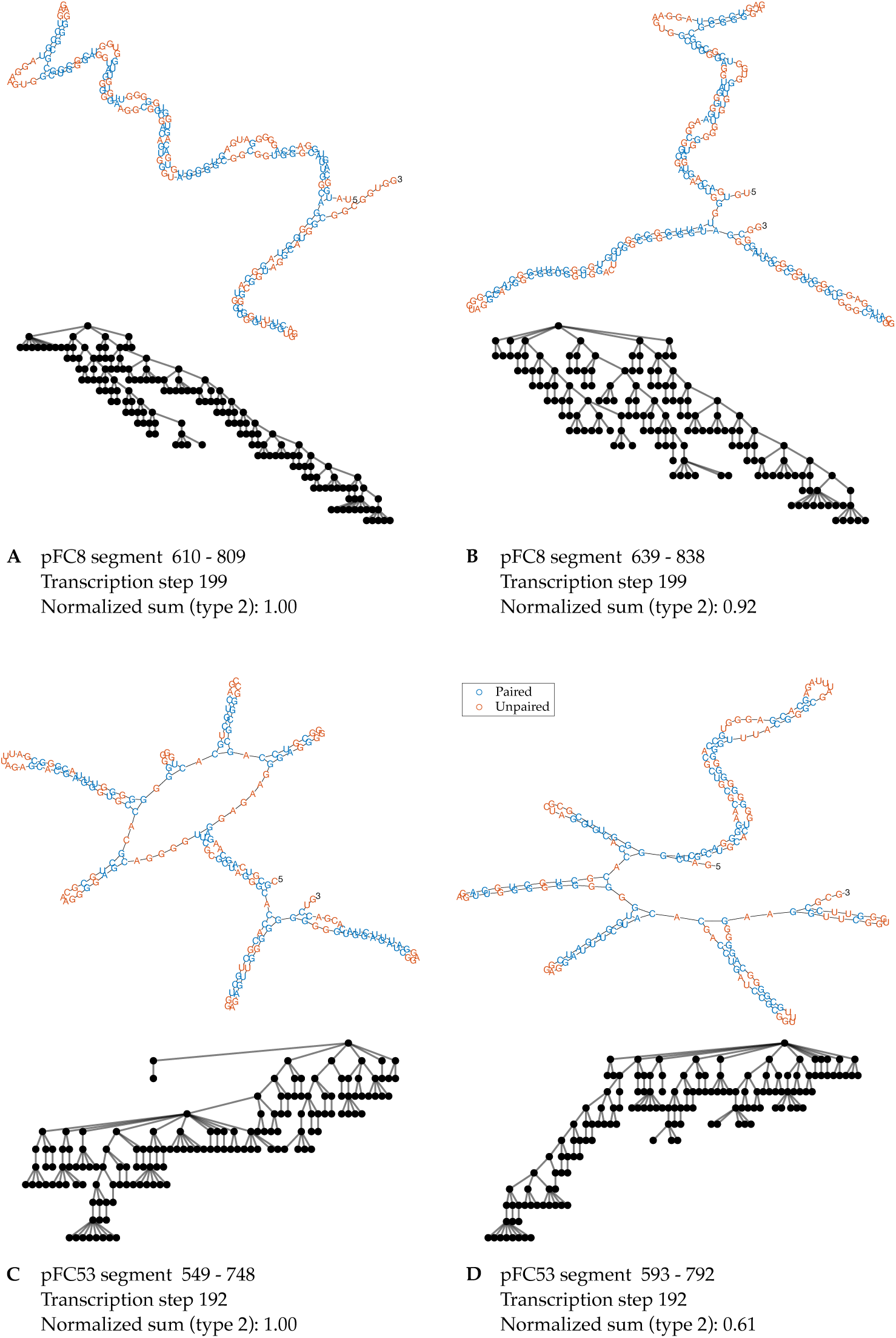
Secondary structures of RNA segments that have the largest type 2 coefficient sums. The figure shows the secondary structures of RNA segments of the pFC8 plasmid (panel A and B) and the pFC53 plasmid (panel C and D) with the two largest type 2 coefficient sums at the transcription step with the highest PCC in the last 10 transcription steps. Type 2 tree representations are displayed following the corresponding secondary structures.

**Supplementary Figure 13:**
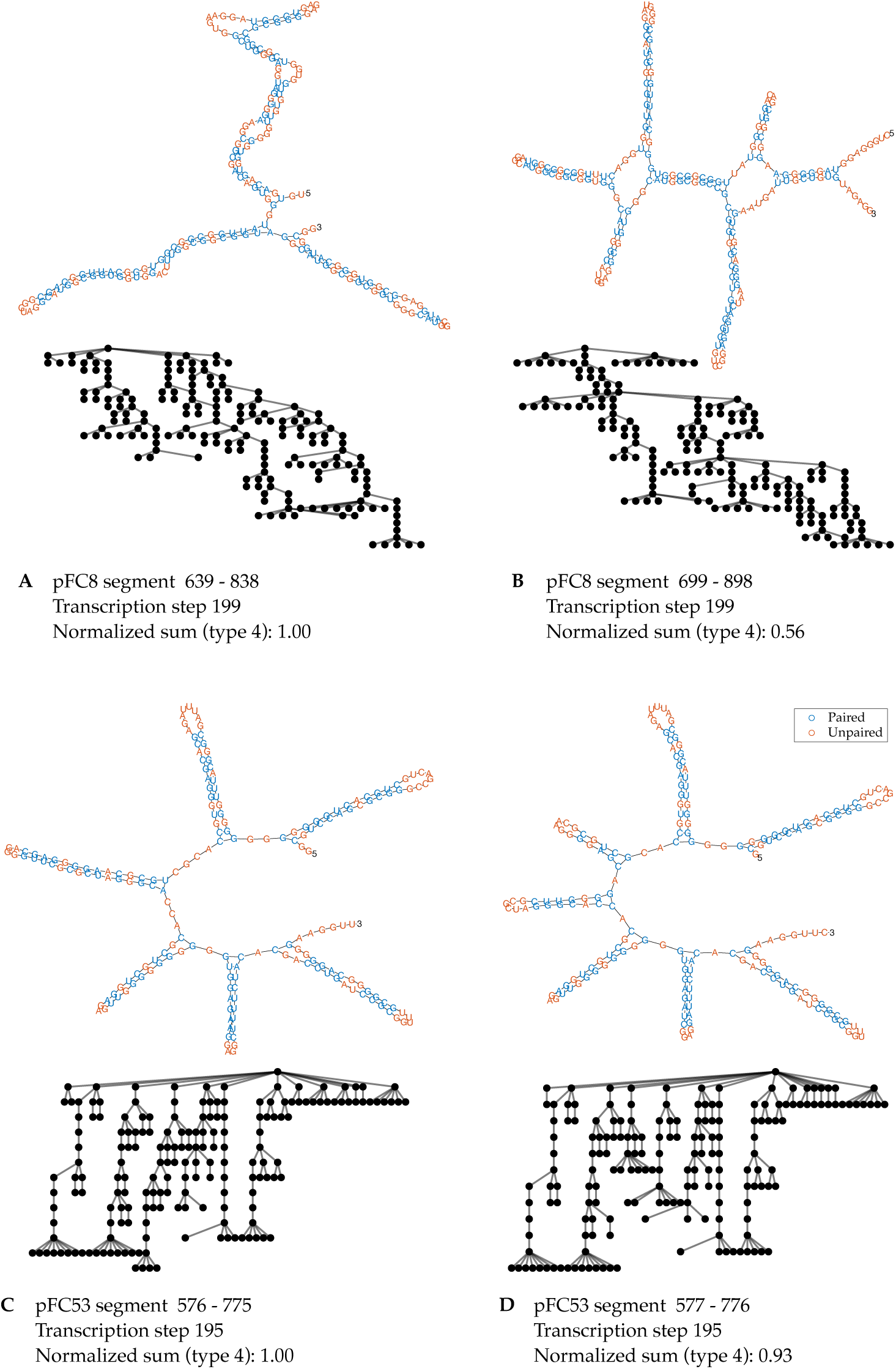
Secondary structures of RNA segments that have the largest type 4 coefficient sums. The figure shows the secondary structures of RNA segments of the pFC8 plasmid (panel A and B) and the pFC53 plasmid (panel C and D) with the two largest type 4 coefficient sums at the transcription step with the highest PCC in the last 10 transcription steps. Type 4 tree representations are displayed following the corresponding secondary structures.

**Supplementary Figure 14:**
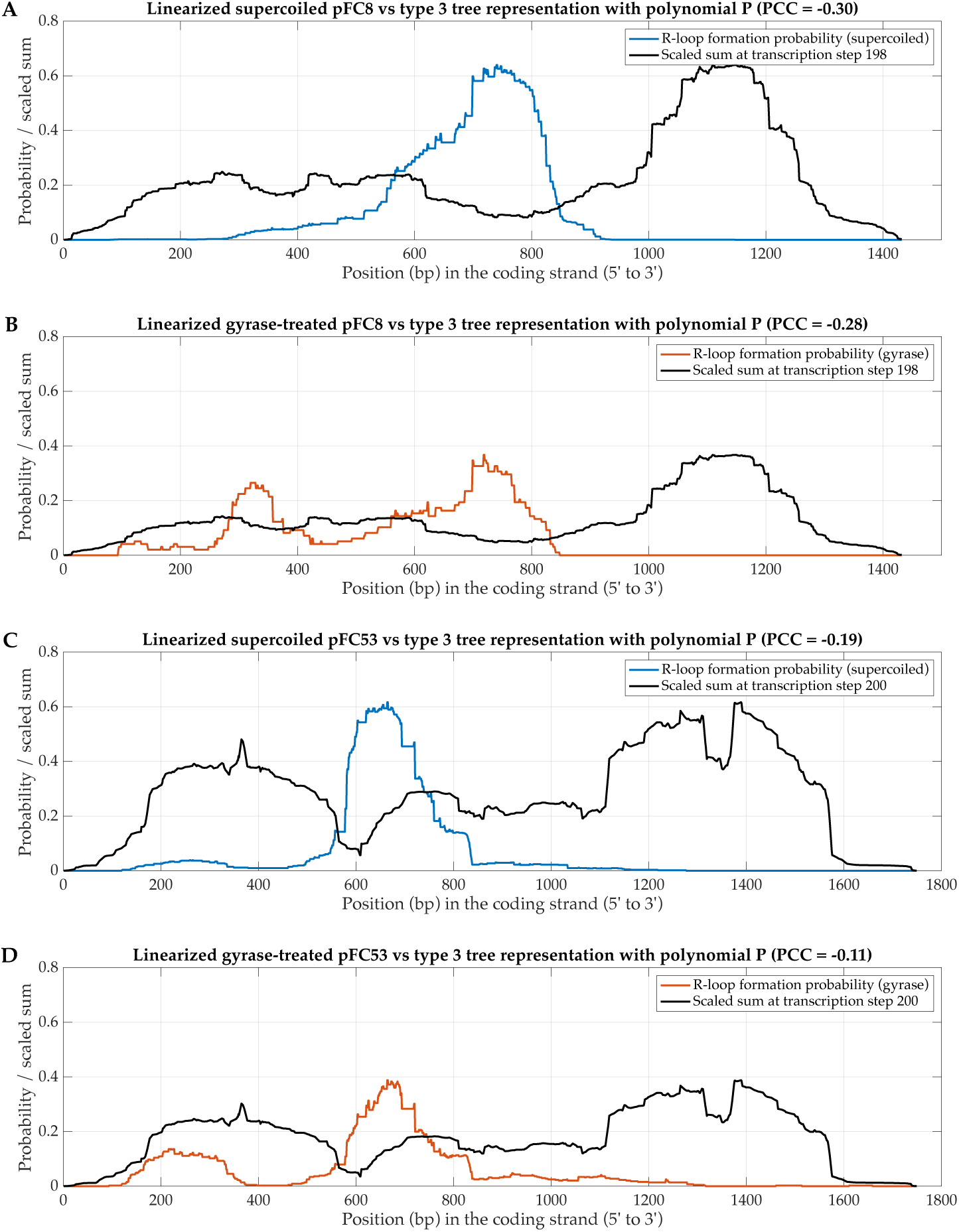
The correlations between the type 3 scaled sums and the R-loop formation probability of pFC8 and pFC53 plasmids. The figure shows the correlations between the type 3 scaled sums (with the highest PCC in the last 10 transcription steps) and the R-loop formation probabilities of the linearized supercoiled pFC8 plasmid (panel A) and of the linearized gyrase-treated pFC8 plasmid (panel B), and the correlations between the type 3 scaled sums (with the highest PCC in the last 10 transcription steps) and the R-loop formation probabilities of the linearized supercoiled pFC53 plasmid (panel C) and of the linearized gyrase-treated pFC53 plasmid (panel D).

**Supplementary Figure 15:**
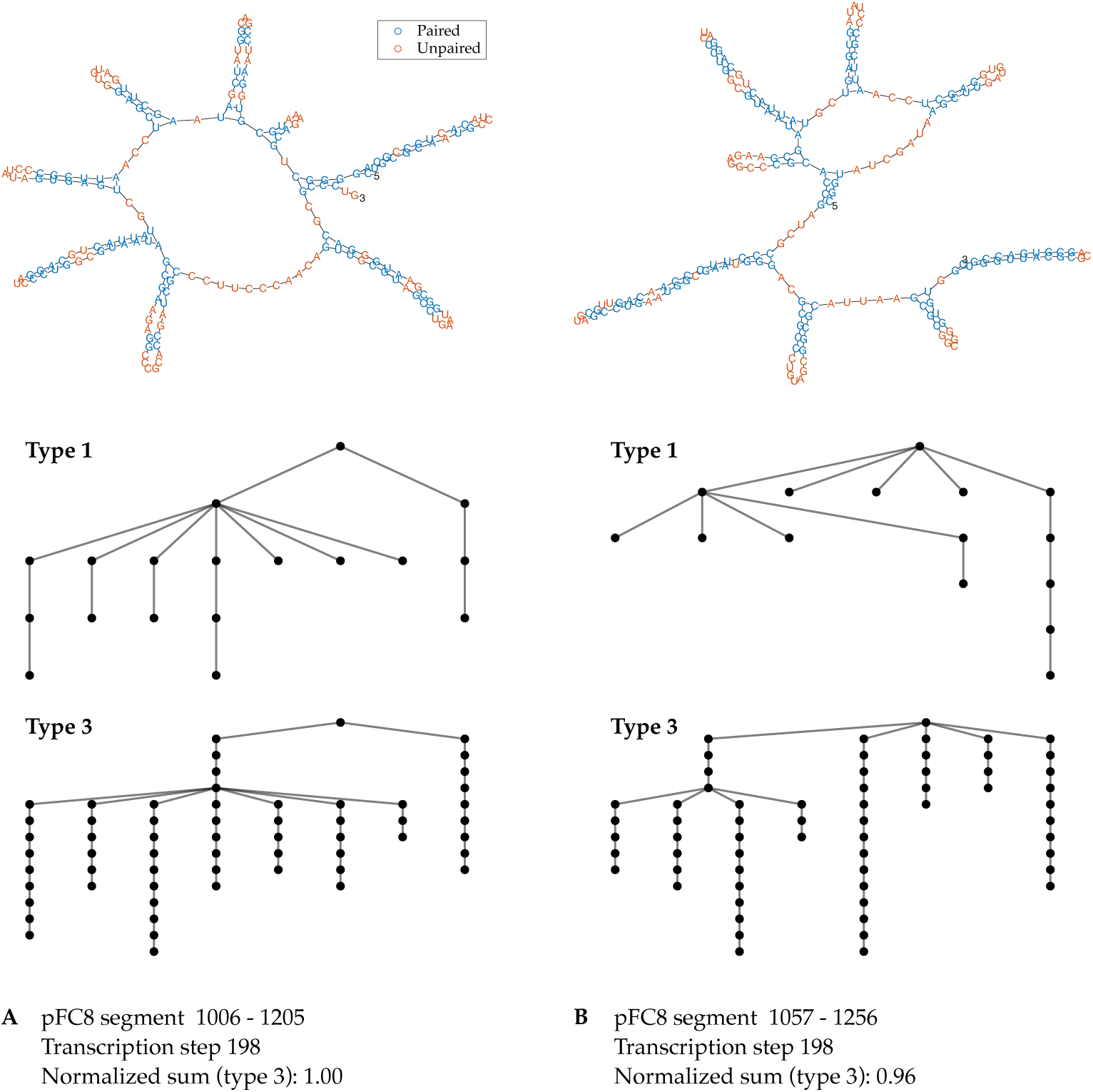
Secondary structures of pFC8 RNA segments that have the largest type 3 coefficient sums. The figure shows the secondary structures of RNA segments of the pFC8 plasmid with the two largest type 3 coefficient sums at the transcription step with the highest PCC in the last 10 transcription steps. Type 1 and type 3 tree representations are displayed following the corresponding secondary structures.

**Supplementary Figure 16:**
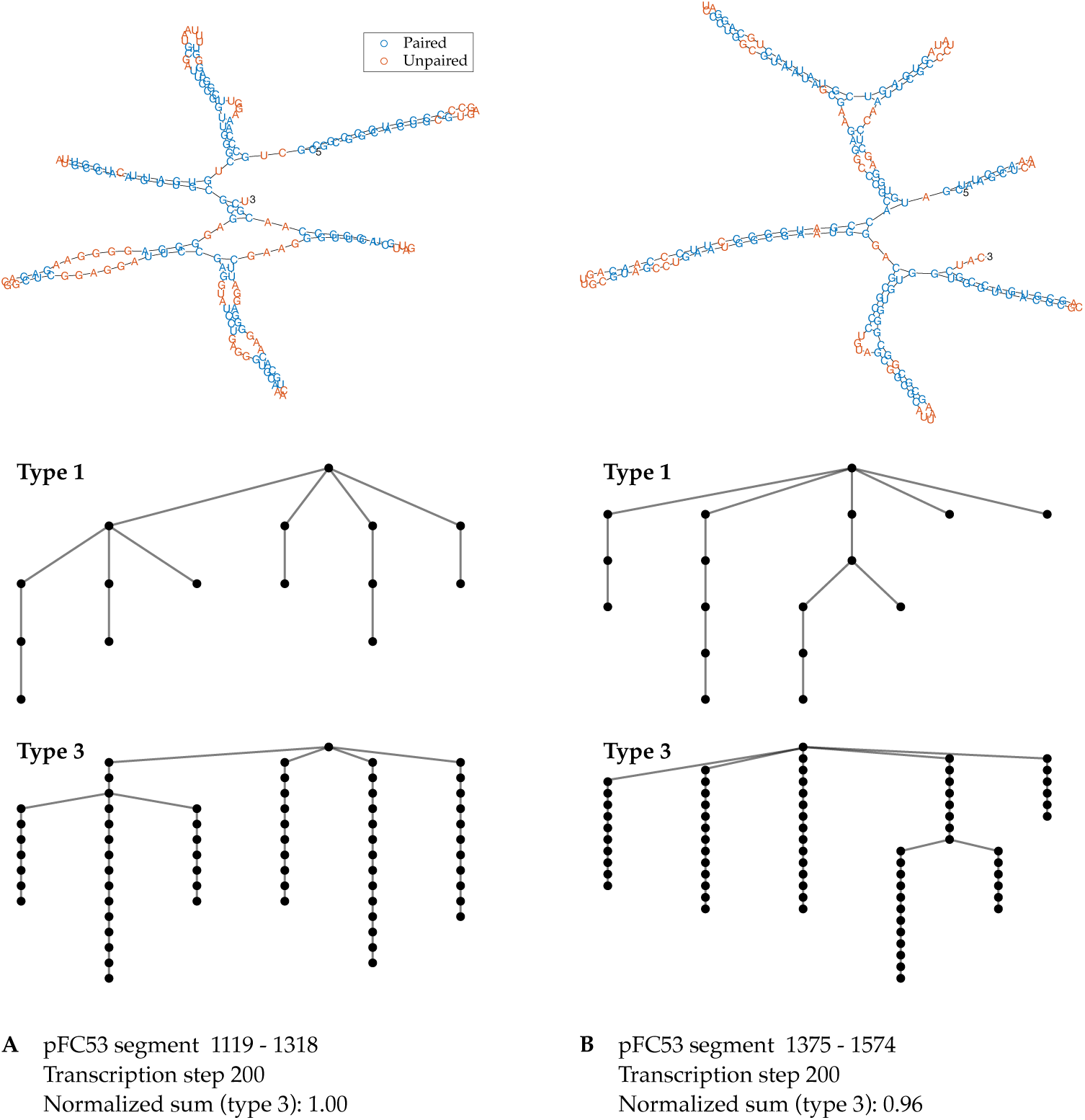
Secondary structures of pFC53 RNA segments that have the largest type 3 coefficient sums. The figure shows the secondary structures of RNA segments of the pFC53 plasmid with the two largest type 3 coefficient sums at the transcription step with the highest PCC in the last 10 transcription steps. Type 1 and type 3 tree representations are displayed following the corresponding secondary structures.

**Supplementary Figure 17:**
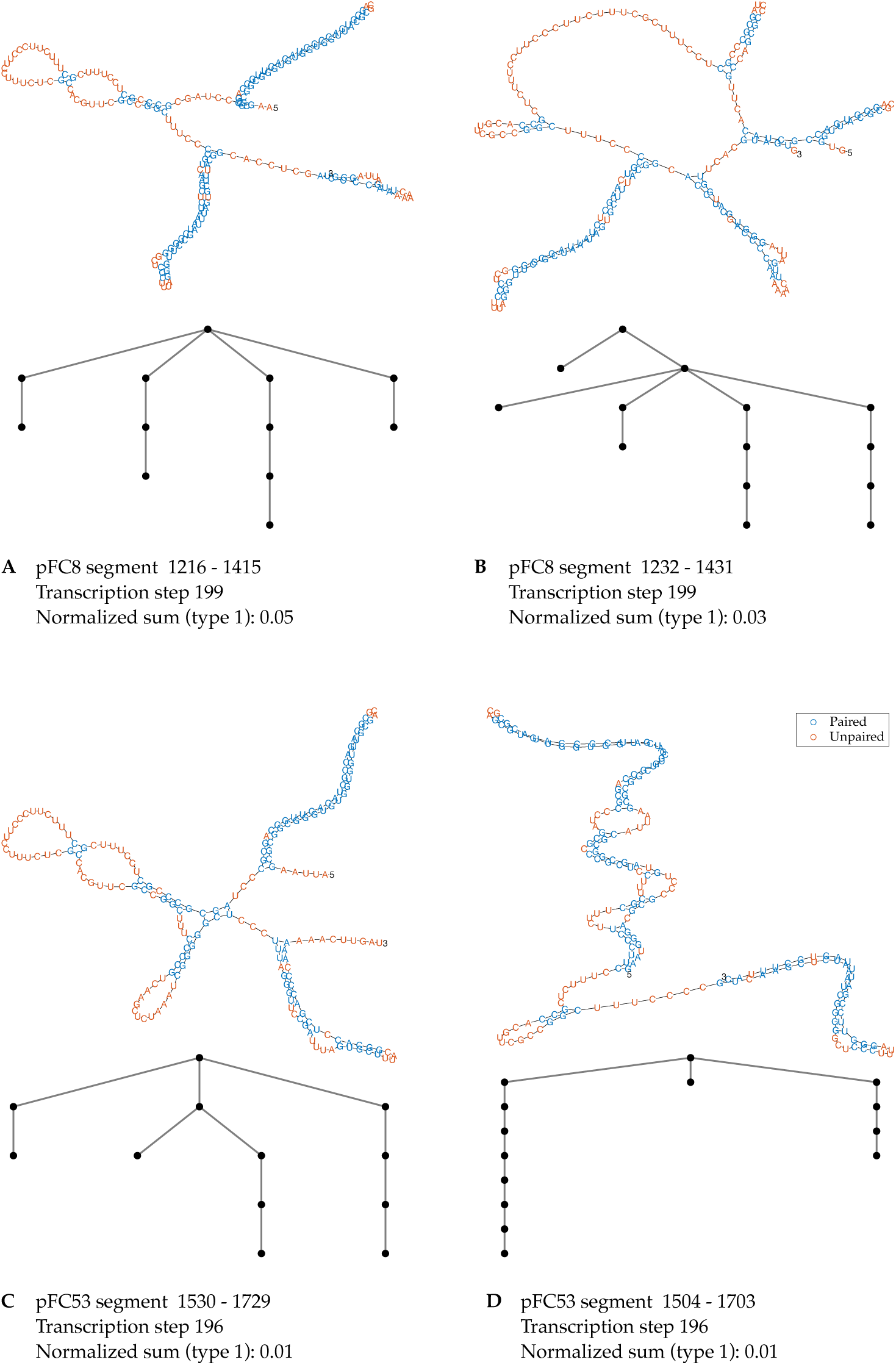
Secondary structures of RNA segments near the 3’ end of the amplicon region. The figure shows the secondary structures of RNA segments of the pFC8 plasmid (panel A and B) and the pFC53 plasmid (panel C and D) near the 3’ end of the amplicon region that have the two largest type 1 coefficient sums at the transcription step with the highest PCC in the last 10 transcription steps. Type 1 tree representations are displayed following the corresponding secondary structures.

**Supplementary Figure 18:**
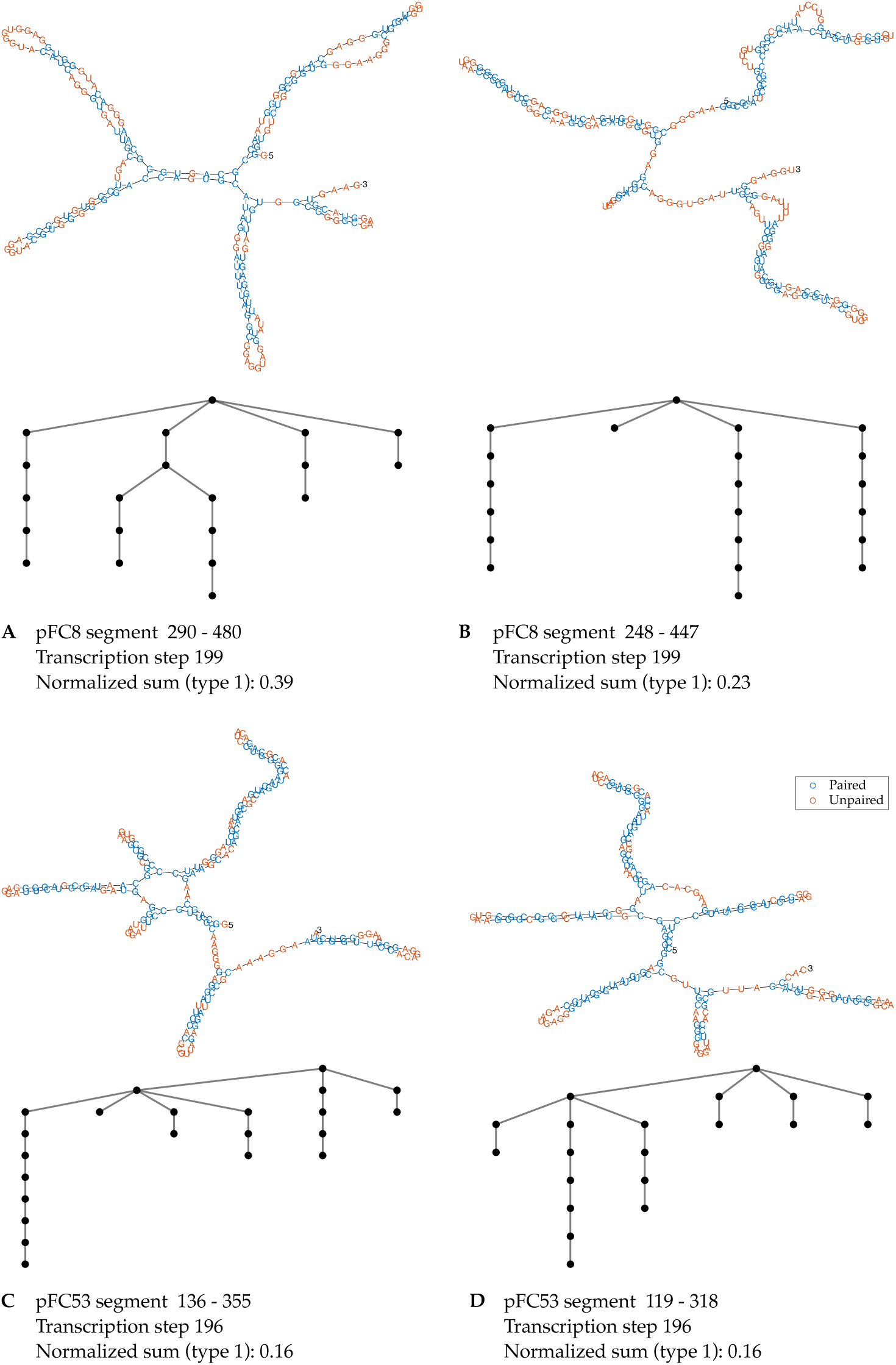
Secondary structures of RNA segments in the minor peak of R-loop formation. The figure shows the secondary structures of RNA segments of the pFC8 plasmid (panel A and B) and the pFC53 plasmid (panel C and D) in the minor peak of R-loop formation that have the two largest type 1 coefficient sums at the transcription step with the highest PCC in the last 10 transcription steps. Type 1 tree representations are displayed following the corresponding secondary structures.

## References

[1] S. Badelt, R. Lorenz, and I. L. Hofacker. DrTransformer: heuristic cotranscriptional RNA folding using the nearest neighbor energy model. Bioinformatics, 39 (1):btad034, 2023.

[2] B. P. Belotserkovskii, S. Tornaletti, A. D. D’Souza, and P. C. Hanawalt. R-loop generation during transcription: Formation, processing and cellular outcomes. DNA Repair, 71:69–81, 2018.

[3] Y. Carrasco-Salas, A. Malapert, S. Sulthana, B. Molcrette, L. Chazot-Franguiadakis, P. Bernard, F. Chédin, C. Faivre-Moskalenko, and V. Vanoost-huyse. The extruded non-template strand determines the architecture of R-loops. Nucleic Acids Research, 47(13):6783–6795, 2019.

[4] F. Chédin. Nascent connections: R-loops and chromatin patterning. Trends in Genetics, 32(12):828–838, 2016.

[5] P. Danaee, M. Rouches, M. Wiley, D. Deng, L. Huang, and D. Hendrix. bpRNA: large-scale automated annotation and analysis of RNA secondary structure. Nucleic Acids Research, 46(11):5381–5394, 2018.

[6] P. A. Ginno, P. L. Lott, H. C. Christensen, I. Korf, and F. Chédin. R-loop formation is a distinctive characteristic of unmethylated human CpG island promoters. Molecular Cell, 45(6):814–825, 2012.

[7] S. Griffiths-Jones, A. Bateman, M. Marshall, A. Khanna, and S. R. Eddy. Rfam: an RNA family database. Nucleic Acids Research, 31(1):439–441, 2003.

[8] S. R. Hartono, A. Malapert, P. Legros, P. Bernard, F. Chédin, and V. Vanoost-huyse. The affinity of the s9.6 antibody for double-stranded rnas impacts the accurate mapping of r-loops in fission yeast. Journal Molecular Biology, 430(3): 272–284, Feb 2018.

[9] Y. Hori, C. Engel, and T. Kobayashi. Regulation of ribosomal RNA gene copy number, transcription and nucleolus organization in eukaryotes. Nature Reviews Molecular Cell Biology, 24(6):414–429, 2023.

[10] R. Janssen and P. Liu. Comparing the topology of phylogenetic network generators. Journal of bioinformatics and computational biology, 19:2140012, 2021.

[11] J. Jedwab, T. Petrie, and S. Simon. An infinite class of unsaturated rooted trees corresponding to designable RNA secondary structures. Theoretical Computer Science, 833:147–163, 2020.

[12] V. F. R. Jones. A polynomial invariant for knots via von neumann algebras. Bulletin of the American Mathematical Society, 12(1):103–111, 1 1985.

[13] P. Liu. A tree distinguishing polynomial. Discrete Applied Mathematics, 288:1–8, 2021.

[14] P. Liu, P. Biller, M. Gould, and C. Colijn. Analyzing phylogenetic trees with a tree lattice coordinate system and a graph polynomial. Systematic Biology, 71(6): 1378–1390, 2022.

[15] P. Liu, T. Feng, and R. Liu. Quantifying syntax similarity with a polynomial representation of dependency trees. Glottometrics, 53:59–79, 2022.

[16] T. J. Macke, D. J. Ecker, R. R. Gutell, D. Gautheret, D. A. Case, and R. Sampath. RNAMotif, an RNA secondary structure definition and search algorithm. Nucleic Acids Research, 29(22):4724–4735, 2001.

[17] M. Malig and F. Chedin. Characterization of R-Loop structures using single-molecule R-Loop footprinting and sequencing. In U. Ørom, editor, RNA-Chromatin Interactions: Methods and Protocols. Springer US, New York, NY, 2020.

[18] M. Malig, S. R. Hartono, J. M. Giafaglione, L. A. Sanz, and F. Chedin. Ultra-deep coverage single-molecule R-loop footprinting reveals principles of R-loop formation. Journal of Molecular Biology, 432(7):2271–2288, 2020.

[19] J. S. Mattick and I. V. Makunin. Non-coding RNA. Human Molecular Genetics, 15: R17–R29, 2006.

[20] J. C. Pons, T. M. Coronado, M. Hendriksen, and A. Francis. A polynomial in-variant for a new class of phylogenetic networks. PLOS ONE, 17(5):1–22, 05 2022.

[21] M. Quadrini, L. Tesei, and E. Merelli. An algebraic language for RNA pseudo-knots comparison. BMC Bioinformatics, 20(4):161, 2019.

[22] J. Santos-Pereira and A. Aguilera. R loops: new modulators of genome dynamics and function. Nature Reviews Genetics, 16(10):583–597, 2015.

[23] L. A. Sanz, S. R. Hartono, Y. W. Lim, S. Steyaert, A. Rajpurkar, P. A. Ginno, X. Xu, and F. Chédin. Prevalent, dynamic, and conserved R-loop structures associate with specific epigenomic signatures in mammals. Molecular Cell, 63(1):167–178, Jul 2016.

[24] S. Schirmer, Y. Ponty, and R. Giegerich. Introduction to RNA secondary structure comparison. Methods in Molecular Biology, 1097:247–273, 2014.

[25] T. Schlick. Adventures with RNA graphs. Methods, 143(1):16–33, 2018.

[26] T. Schlick, Q. Zhu, A. Dey, S. Jain, S. Yan, and A. Laederach. To knot or not to knot: Multiple conformations of the SARS-CoV-2 frameshifting RNA element. Journal of the American Chemical Society, 143(30):11404–11422, 08 2021.

[27] W. R. Schmitt and M. S. Waterman. Linear trees and RNA secondary structure. Discrete Applied Mathematics, 51(3):317–323, 1994.

[28] E. Schubert and P. J. Rousseeuw. Fast and eager k-medoids clustering: O(k) runtime improvement of the PAM, CLARA, and CLARANS algorithms. Information Systems, 101:101804, 2021.

[29] R. Stolz, S. Sulthana, S. R. Hartono, M. Malig, C. J. Benham, and F. Chedin. Interplay between DNA sequence and negative superhelicity drives R-loop structures. Proceedings of the National Academy of Sciences, 116(13):6260–6269, 2019.

[30] W. T. Tutte. A contribution to the theory of chromatic polynomials. Canadian Journal of Mathematics, 6:80–91, 1954.

[31] L. van Iersel, V. Moulton, and Y. Murakami. Polynomial invariants for cactuses. Information Processing Letters, 182:106394, 2023.

[32] W. Xu, H. Xu, K. Li, Y. Fan, Y. Liu, X. Yang, and Q. Sun. The R-loop is a common chromatin feature of the Arabidopsis genome. Nature Plants, 3(9):704–714, 2017.

